# Interpretable machine learning models for single-cell ChIP-seq imputation

**DOI:** 10.1101/2019.12.20.883983

**Authors:** Steffen Albrecht, Tommaso Andreani, Miguel A. Andrade-Navarro, Jean-Fred Fontaine

## Abstract

**Motivation:** Single-cell ChIP-seq (scChIP-seq) analysis is challenging due to data sparsity. High degree of data sparsity in biological high-throughput single-cell data is generally handled with imputation methods that complete the data, but specific methods for scChIP-seq are lacking. We present SIMPA, a scChIP-seq data imputation method leveraging predictive information within bulk data from ENCODE to impute missing protein-DNA interacting regions of target histone marks or transcription factors.

**Results:** Imputations using machine learning models trained for each single cell, each target, and each genomic region accurately preserve cell type clustering and improve pathway-related gene identification on real data. Results on simulated data show that 100 input genomic regions are already enough to train single-cell specific models for the imputation of thousands of undetected regions. Furthermore, SIMPA enables the interpretation of machine learning models by revealing interaction sites of a given single cell that are most important for the imputation model trained for a specific genomic region. The corresponding feature importance values derived from promoter-interaction profiles of H3K4me3, an activating histone mark, highly correlate with co-expression of genes that are present within the cell-type specific pathways. An imputation method that allows the interpretation of the underlying models facilitates users to gain an even deeper understanding of individual cells and, consequently, of sparse scChIP-seq datasets.

**Availability and implementation:** Our interpretable imputation algorithm was implemented in Python and is available at https://github.com/salbrec/SIMPA

## Introduction

The discovery of protein-DNA interactions of histone marks and transcription factors is of great importance in biomedical studies because of their impact on the regulation of core cellular processes such as chromatin structure organization and gene expression. These interactions are measured by chromatin immunoprecipitation followed by high-throughput sequencing (ChIP-seq). Public data from the ENCODE portal, which provides a large collection of experimental bulk ChIP-seq data, has been used for comprehensive investigations providing insights into epigenomic processes that affect chromatin 3D-structure, chromatin state, and gene expression, to name just a few (Consortium and others, 2012).

Recently developed protocols for scChIP-seq are powerful techniques that will enable in-depth characterization of those processes at single-cell resolution. ChIP-seq was successfully performed on single cells with sequencing depth as low as 1,000 unique reads per cell, reflecting the low amount of cellular material that can be obtained from only one single cell (Rotem *et al*., 2015). Even though this low coverage leads to sparse datasets, scChIP-seq data has enabled the study of biological systems that cannot be investigated with bulk ChIP-seq applied for millions of cells, for example, the differences between drug-sensitive and drug-resistant breast cancer cells (Grosselin *et al*., 2019).

Nevertheless, the analysis of single-cell assays is strongly affected by the sparsity of data. In the context of ChIP-seq, sparsity means no signal observed for numerous genomic regions without the possibility to explain whether this is real or due to low sequencing coverage. Notably, sparsity may disable the investigation of functional genomic elements that could be of crucial interest. Hence, an imputation method is needed to complete sparse scChIP-seq datasets while preserving the identity of each individual cell.

The first published imputation method for NGS epigenomic signals was ChromImpute (Ernst and Kellis, 2015), later followed by (Durham *et al*., 2018), an improved method for the imputation of signal tracks for several molecular assays in a biosample-specific manner (*biosample* refers to the specific tissue or cell-type). The challenge of transcription factor binding site prediction was approached, for example, using deep learning algorithms on sequence position weight matrices (Qin and Feng, 2017), and more recently by the embedding of transcription factor labels and k-mers (Yuan *et al*., 2019). With the aim to complete the ENCODE portal with imputed bulk experiments, Schreiber *et al*. implemented the method Avocado, which extends the basic concept of PREDICTD by deep neural networks (Schreiber, Durham, *et al*., 2020). Avocado was also validated on ChIP-seq data from both histone marks and transcription factors (Schreiber, Bilmes and Noble, 2020). Such methods show the successful application of machine learning algorithms and mathematical approaches in predicting epigenomic signals such as transcription factor binding activity. However, their scope, being limited to either imputation of missing bulk experiments or sequence-specific binding site prediction, hampers their application to single-cell data.

The challenge of imputation for sparse datasets from single-cell assays has been extensively approached for single-cell RNA-seq (scRNA-seq) used to quantify gene expression at single-cell resolution (Ronen and Akalin, 2018; Zhang and Zhang, 2018; Peng *et al*., 2019; Chen *et al*., 2020; Elyanow *et al*., 2020; Jeong and Liu, 2020; Tang *et al*., 2020; Ye *et al*., 2020; Zhu and Anastassiou, 2020). In this context, similarly to scChIP-seq data, sparsity is described by dropout events, which are transcripts having a transcription rate of zero without knowing if the corresponding gene is not expressed at all or if the expression rate is not detected due to technical limitations (Tang *et al*., 2020). The question arises if these methods can be easily adapted for imputation of scChIP-seq data. However, there are crucial differences between the application of RNA-seq and ChIP-seq techniques that must be considered regarding the development of a method for scChIP-seq imputation.

First, in RNA-seq the set of relevant genomic regions, defined by the species-specific transcripts, is more limited. For a ChIP-seq profile, the regions of potential interest may originate from any position in the genome and cannot be defined in advance. To simplify the analysis, in scChIP-seq imputation the genome can be organized in non-overlapping genomic windows (bins) of a certain size. At 5 kb resolution, this binning concept results in more than 600,000 possible regions in the human genome, a number that is much higher than the number of transcripts considered in the RNA-seq context.

The second main difference is that scChIP-seq interactions are usually represented by a Boolean value describing the presence or absence of a significant enrichment of sequencing reads defining a peak, while RNA-seq datasets contain transcription rates. Consequently, the application of scRNA-seq imputation methods on scChIP-seq data might be less appropriate.

In contrast, imputation methods for chromatin accessibility profiles from single-cell ATAC-seq (single-cell Assay for Transposase-Accessible Chromatin using sequencing, scATAC-seq) are potentially more transferable to scChIP-seq imputation as their data representation is more similar. A few methods exist that implement imputation for scATAC-seq, though none of them was tested on scChIP-seq data so far. Methods such as SCALE (Xiong *et al*., 2019), FITs (Sharma *et al*., 2020) and scOpen (Z. Li *et al*., 2019) have been shown to outperform scRNA-seq methods to impute scATAC-seq data. These methods complement each other with respect to the different approaches they implement, however, they share the common concept of imputing the missing values within a sparse matrix defined by the single cells (rows) and genomic bins (columns), and only bins are considered that were detected by at least one single cell if no further filtering is applied. Consequently, such methods can offer imputation only on regions that were observed in the single-cell dataset and it is likely that many important regions along the whole genome will be missed.

To overcome this limitation, we developed SIMPA, an algorithm for **S**ingle-cell Ch**I**P-seq i**MP**ut**A**tion, that uses bulk ChIP-seq datasets of the ENCODE project (“The ENCODE (ENCyclopedia Of DNA Elements) Project,” 2004; Sloan *et al*., 2016). It was already shown that an additional bulk RNA-seq dataset can be used to improve the imputation for a sparse scRNA-seq dataset (Peng *et al*., 2019). Within SIMPA, publicly available bulk ChIP-seq data is turned into a reference set used to define potential bins to be imputed and then leveraged by machine learning models to compute specific imputation probabilities. Moreover, these models are interpretable and can be used to gain more insights into a given single-cell dataset, allowing the investigation of individual cells on a more detailed level. The interpretability is implemented by InterSIMPA, an extension of SIMPA, which takes a single cell as input together with a genomic position of interest and trains one classification model for the position to derive a probability which can be seen as an imputation score. More importantly, InterSIMPA ranks the genomic regions from the sparse single cell profile by their relevance for the model. The ranked regions are enriched by detailed information and an importance score describing the strength of their relationship with the given genomic position of interest. These relationships can be interpreted as dependencies between genomic regions that could be part of the gene regulatory network (e.g., between enhancers and promoters).

The basic reference-based imputation concept of SIMPA was first validated on simulated data and then on a real scChIP-seq dataset of the H3K4me3 and H3K27me3 histone marks in B-cells and T-cells (Grosselin *et al*., 2019). The latter dataset allowed us to investigate the algorithm’s capability of retaining the cell-type clustering and furthermore to assess the biological relevance of the imputed regions based on a pathway enrichment analysis. These validations were performed in comparison to a reference-free method represented by the aforementioned scATAC-seq analysis method SCALE, which applies a combination of Gaussian mixture models and variational autoencoder (Xiong *et al*., 2019). Results from InterSIMPA were validated using promoter regions related to genes of the B- and T-cell signaling pathways, and compared to gene co-expression data derived from the STRING database (Szklarczyk *et al*., 2019).

## Results

### Algorithmic concept and cross-validations

Unlike many other single-cell imputation methods, SIMPA leverages predictive information within bulk ChIP-seq data by combining the sparse input of one single cell and a collection of 2,251 ChIP-seq experiments from ENCODE. In order to better compare bulk and single-cell data, ChIP-seq regions (or significant signal/noise ChIP-seq peaks) are mapped to genomic bins (**Fig. 1A** and **Supplementary Note 2; and Supplementary Note 1 for details about bulk and single-cell data retrieval and processing**).

**Figure 1.**
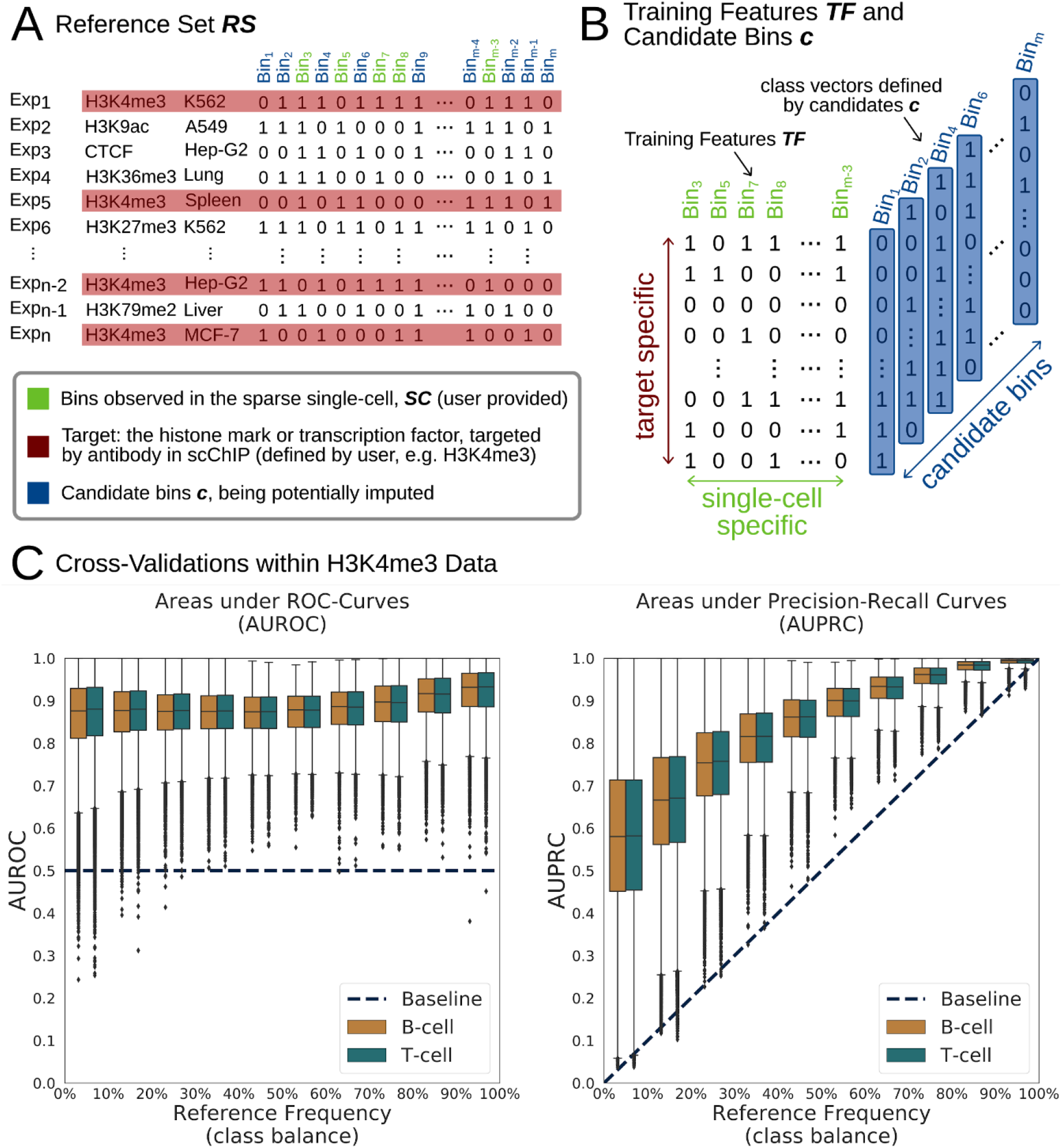
SIMPA’s algorithm and cross validations. **A**. Identified ChIP-seq regions from bulk experiments were downloaded from ENCODE and mapped to bins defined as non-overlapping and contiguous genomic regions of a defined length (5 kb for H3K4me3 and 50 kb for H3K27me3) and covering the whole genome (the table). A bin is given a value of 1 for a particular experiment if there is at least one ChIP-seq region in this experiment that overlaps the bin, 0 otherwise. In total 2251 ChIP-seq experiments for several targets (histone marks or transcription factors) performed in several biosamples (tissues and cell-lines) were downloaded from ENCODE portal and preprocessed. Depending on the target specified by the user, the target-specific reference set **RS** is then created and contains all experiments related to this target (red lines) and all bins observed for at least one of those experiments. **B**. The single-cell specific training feature matrix **TF** is created as a subset of **RS** by selecting only bins observed within the given single cell (green columns). All other bins from **RS** are the candidate bins (**c**; blue columns) and define the class vectors consisting of the corresponding values in **RS**. For each candidate bin, a classification model is trained based on the training features and the class vector identifying associated experiments. **C**. Cross-validated evaluations of SIMPA’s Random Forest performances to predict values of candidate bins in single cells within the H3K4me3 data. For each bin, a ten-fold cross-validation was applied and summarized as Area under ROC-Curve (AUROC) or Area under Precision-Recall Curve (AUPRC) (y-axes). Results for all bins are represented by boxplots subdivided by class balance in the candidate bins (percentage of “1” values in the bin) (x-axis). The dashed lines describe the baseline performance expected from a random classification model: 0.5 for AUROC and equal to the class balance for AUPRC.

SIMPA produces results for each single cell of a scChIP-seq dataset by using machine learning models trained on a subset of the ENCODE data related to a selected target, that is the histone mark or transcription factor used in the single-cell experiment. Derived from this target-specific subset, the classification features are defined by genomic regions detected in the single cell, while the class to predict is defined by a region observed in at least one target-specific bulk ENCODE experiment, but not in the single cell (**Fig. 1B**). In other words, by using this particular data selection strategy, SIMPA searches relevant statistical patterns linking (i) protein-DNA interacting regions across single-cell specific regions of the target-specific ENCODE data for different cell types to (ii) the presence or absence of a potential region for the given single cell. SIMPA’s machine learning models are able to use those patterns to provide accurate predictions (**Fig. 1C, Supplementary Note 3**, and **Fig. S1**). Moreover, on the high-resolution H3K4me3 dataset, SIMPA achieved high recall rates for bins removed from single-cell profiles (**Supplementary Note 4** and **Fig. S2**). The single-cell profiles used to create the training feature sets within the cross-validations were defined by the dataset from Grosselin *et al*. (Grosselin *et al*., 2019) called the real scCHIP-seq dataset below.

### Validation on simulated data

In order to evaluate the algorithm’s ability from few input bins (hundreds) to complete full data profiles (thousands of bins) of different protein targets and cell-types, we simulated sparse protein-DNA interaction profiles from the bulk ENCODE ChIP-seq experiments that are used as reference data by SIMPA. For the simulation, we took bulk experiments for different cell-type-target combinations to define them as full single-cell profiles (*origin*) and down-sampled those profiles to simulate sparse single-cell profiles (from 100 to 1600 bins) (see **Supplementary Note 5** for details). Each simulated sparse profile was used as input for SIMPA and the output was compared to the origin. For the model training, the full origin profile was excluded from the reference training set in order to apply the default validation, called *leave-out origin* (LOO). Additionally, a more challenging validation strategy was applied in which all reference profiles for the same cell-type (biosample) were excluded, called *leave-out cell-type* (LOCT).

For H3K4me3, the most frequently investigated target in ENCODE, high area under ROC-curve values confirm that SIMPA is able to accurately recapitulate the original data from the simulated sparse profiles (**Fig. 2A**). Even if the cell-type-specific information is completely removed from the training set (LOCT), the performance is still high. Furthermore, these observations are confirmed when using precision-recall curves as performance measure (**Fig. 2B**), a highly relevant analysis given the imbalance in the validation sets (containing far fewer positive than negative samples). We made similar observations in a ROC-curve and precision-recall curve analysis for other cell-type-target combinations (**Fig. S4 and Fig. S5**).

**Figure 2.**
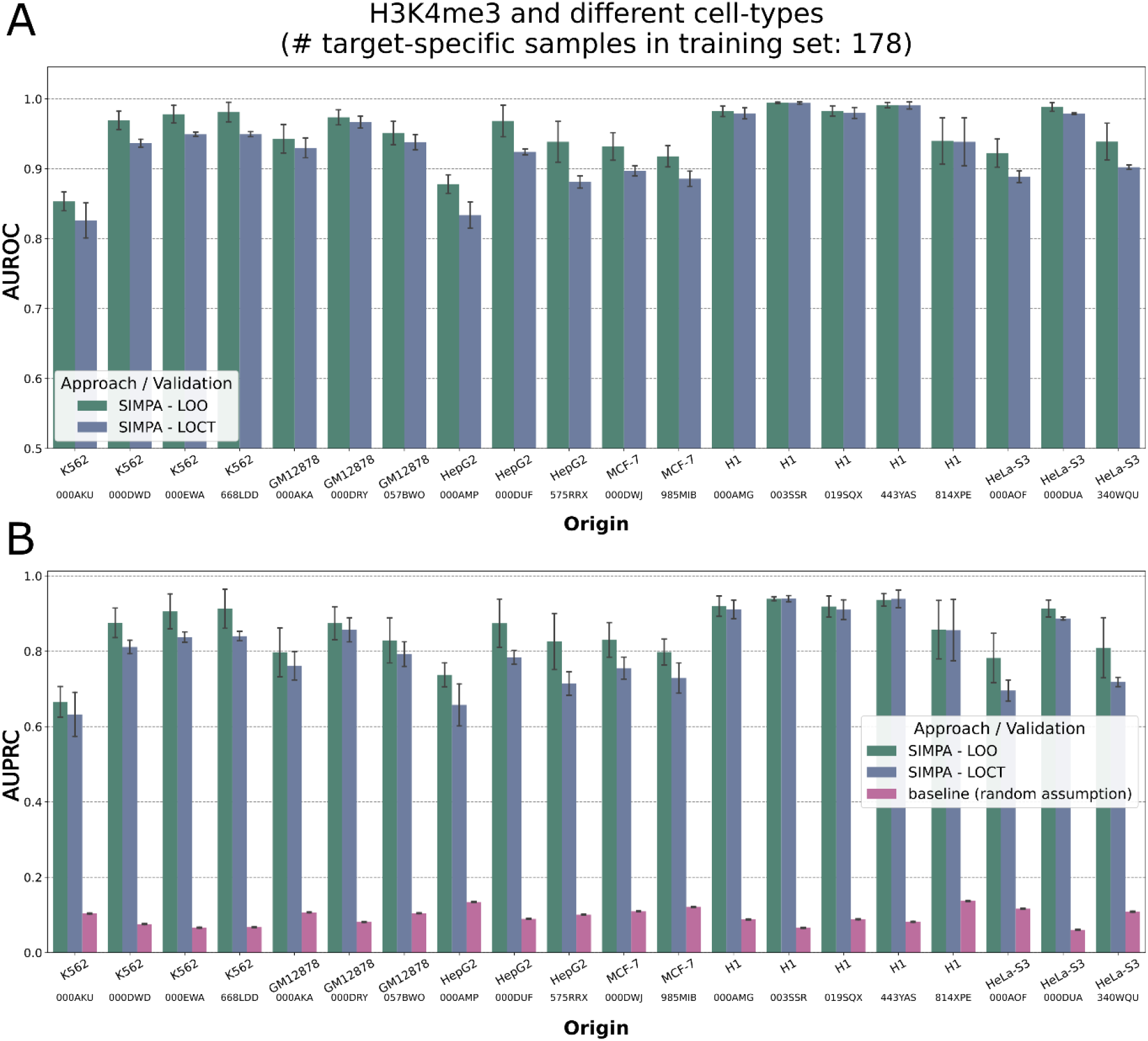
Performance on simulated sparse profiles in different cell-types. **A**. Using the origin profile for simulation as the validation set of true binding interactions, the area under ROC-curve (AUROC in y-axis) describes the capability of SIMPA to accurately impute and recapitulate the bulk experiment. The bars describe the mean AUROC and the error bars describe the standard deviation across multiple applications on sparse sets with different sizes. SIMPA was validated with two strategies, the default leave-out origin (LOO) and the extreme leave-out cell-type (LOCT). The x-axis labels indicate the cell-type of the origin profile and additionally the ENCODE accession to show which of the experimental dataset was used as origin. **B**. Same as in **A** but using the area under precision-recall curve (AUPRC in y-axis) as performance measure. The pink bars show the class balance (fraction of positives in the class feature) representing the random assumption as baseline to be expected from a primitive classifier that randomly assigns the class values (according to (Saito Takaya AND Rehmsmeier, 2015)).

In order to assess the single-cell specificity of SIMPA in this simulation, we compared each fully imputed profile to its origin profile and also to a consensus profile representing experimental datasets that are most similar to the origin (experimental profiles with same protein target and same cell-type, see **Supplementary Note 5** for details). Results show that for most of the simulations (>95%) the imputed profile is closer to the origin profile, hence single-cell specific (**Fig. 3A**). Moreover, we observed that the origin profiles can be more similar to the consensus profile (less specific) or less similar (more specific). When the origin profiles are less specific, it is harder for SIMPA to achieve an imputed profile specific to the origin (single-cell specific). However, for such cases in which the origin is quite close to the consensus (Jaccard-Index > 0.65) the imputation is still single-cell specific, although with a lower single-cell specificity value (**Fig. 3B**).

**Figure 3.**
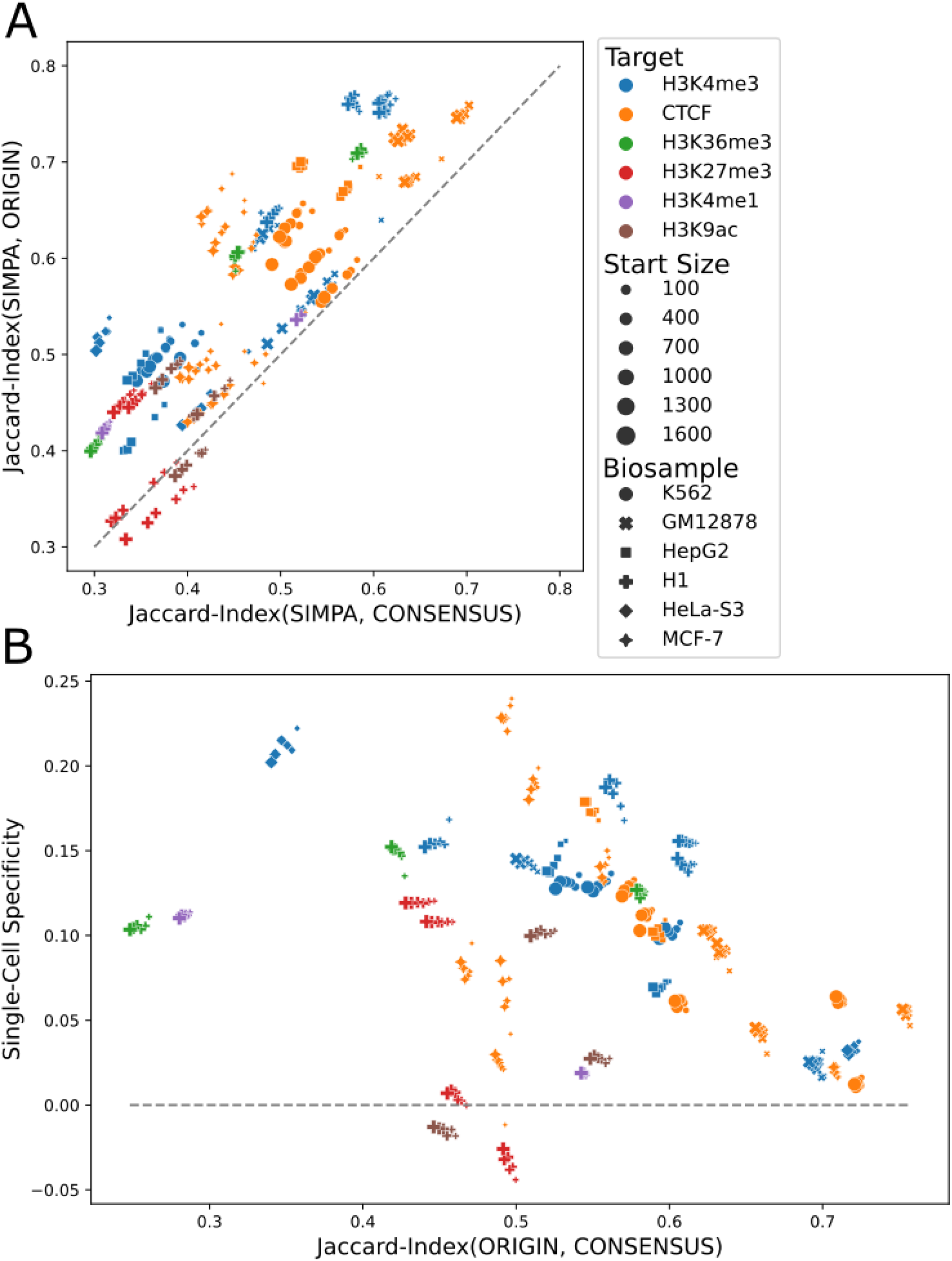
Single-cell specificity analysis. **A**. The Jaccard-Index is used to compare the imputed profiles from SIMPA with the origin profile used to create a simulated sparse profile and the consensus profile representing the remaining experiments available for the same biosample-target combination as the origin profile. The dashed line shows the balance line at which the imputed profile from SIMPA is neither closer to the origin nor to the consensus. Cases above the dashed line are those in which the imputed profile is single-cell specific, hence, closer to the origin than to the consensus. **B**. “Single-Cell Specificity” on the y-axis is defined as the difference between the imputed-to-origin similarity (y-axis in **A**) and imputed-to-consensus similarity (x-axis in **A**). Having the similarity between the origin and the consensus on the x-axis, this plot allows the visualization of the single-cell specificity in relation to how specific the origin is. The higher the similarity between the origin and consensus, the less specific is the origin profile and the harder the challenge to capture its specificity. Profiles, above the 0 line, are single-cell specific as they are closer to the origin than to the consensus.

Taken together, the simulation results show that models trained from a few bins accurately impute thousands of bins and show that completed profiles can be single-cell specific on real data even if the investigated cell-type is not represented by any of the bulk datasets in the reference set (leave-out cell-type validation).

### Model interpretability on real data

Addressing one main aims of this study: to make models interpretable, we implemented an extension called InterSIMPA. Here we define interpretability as the possibility of obtaining information of potential biological relevance from the relationships observed between the training features (genomic regions) and an imputed region of interest. These relations can be expected to be part of the genomic regulatory network.

The training features are derived by InterSIMPA in the same way as for SIMPA but a single machine learning model is trained for a genomic position of interest defined by the user. Accordingly, one imputed probability is returned with information about the genomic regions from the single cell most important for the machine learning model. Finally, the algorithm reports the genes closest to these regions (**Supplementary Note 6**).

To demonstrate how interpretable imputation models can be used to expose more information from the sparse ChIP-seq profile of individual single cells, we use the single-cell ChIP-seq dataset of H3K4me3 interactions in B-cells and T-cells from Grosselin *et al*. According to the given cell types, we focused on promoter regions of genes that are involved within the B-cell and T-cell receptor signaling pathways. The two gene sets contain 67 and 97 genes for the B-cell receptor and T-cell receptor signaling pathways, respectively, with an overlap of 44 genes. To focus on the genes that could be more specific to the cell-types under investigation, from the union of the two gene sets we selected 24 genes with frequency of their promoter regions lower than 20% in the corresponding H3K4me3-specific reference set, which means that their promoter has no detected interaction site for more than 80% of the ENCODE reference experiments for H3K4me3 in different cell-types and tissues (**Supplementary Note 6**).

As H3K4me3 is an activating histone mark, we expected to observe interaction sites in the promoter regions of these genes. However, for many of those promoter regions, the H3K4me3 binding is missing for most of the single cells in the sparse data (**Fig. 4A**). Our expectation that SIMPA is able to impute such regions in a cell-type-specific manner, is confirmed by comparing the imputed probabilities calculated by SIMPA for promoter regions of the 24 selected genes in single B- and T-cells (**Fig. 4B)**. For most of the genes, the imputed probability is higher when SIMPA is applied on single cells that are from the pathway-related cell-type. Finally, we evaluated the interpretability of the 24 imputation models by comparing the feature importance values of the extracted features from the single cell and co-expression values of the feature-related genes with the gene of the imputed promoter (**Fig. 4C**). Co-expression data from the STRING database was used (Szklarczyk *et al*., 2019). The observed high correlations suggest that InterSIMPA is capable to describe biologically relevant promoter-promoter relations by the predictive information hidden within sparse histone mark profiles of an activating mark. Consequently, our approach not only completes the sparse scChIP-seq dataset, but its interpretability-extension is even capable of providing deeper insights into the data.

**Figure 4.**
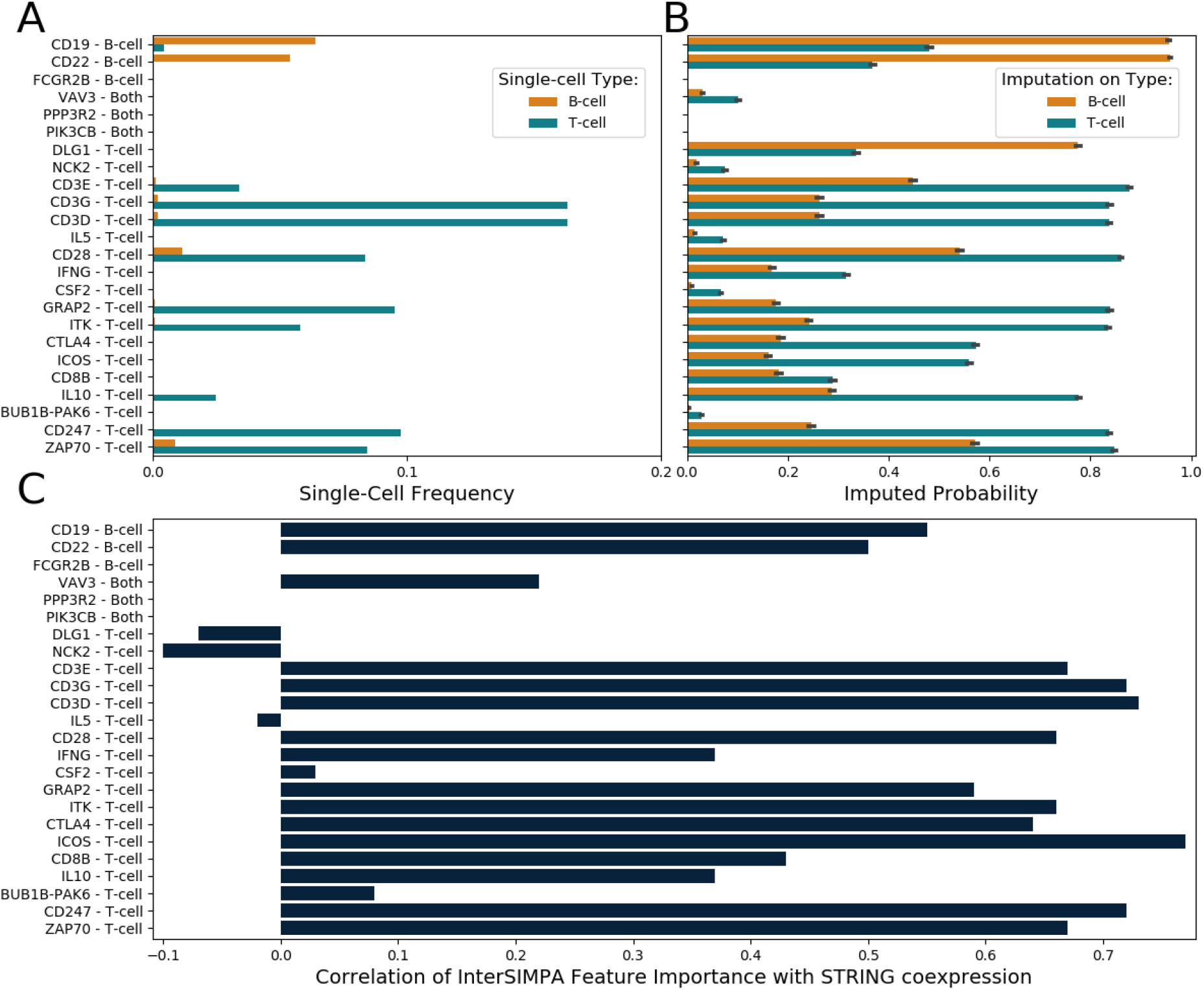
Pathway related gene analysis using the interpretation of imputation models. **A**. Fraction of single cells for which H3K4me3 binding is observed within the gene’s promoter region in the single-cell dataset (orange and blue bars representing B-cells or T-cells, respectively). y-axis labels show the gene names and if the gene belongs to the B-cell or T-cell receptor signaling pathway or to both. **B**. Imputed probability computed by SIMPA for the gene-related promoter regions shown in A. The imputation was applied on numerous single cells from the cell-types B-cell (orange) and T-cell (blue). The error bars represent the standard deviation across the imputation runs on different cells. For the majority of genes, the imputed probability is higher within the cell-type that corresponds to the gene’s pathway. **C**. Correlation of feature importance and co-expression values. For each model used to impute a promoter (y-axis), the training features (genomic bins) were extracted together with their importance value provided by the Random Forest algorithm and annotated with the nearest gene on the genome. Co-expression values, derived from transcriptomic and proteomic measurements, of those genes with the gene related to the imputed promoter were retrieved from the STRING database. The Pearson correlation coefficient of feature importance and co-expression values is shown (x-axis).

### Performance on cell-type clustering and functional analysis

After the evaluation of the InterSIMPA extension, here, we evaluate how SIMPA enhances single cell data corresponding to different cell types. For the imputation of a full single-cell dataset, SIMPA was applied for each cell individually. The resulting imputed profiles were then analyzed within two validations, to examine if (i) cell-type clustering was retained after imputation and (ii) if the imputed single-cell profiles are associated with genes of the corresponding cell-type-specific pathway. Following the investigations of Schreiber *et al*. (Schreiber, Singh, *et al*., 2020), we also compared bin probabilities from SIMPA to a simple imputation approach that uses bin frequencies in the reference set (experiments with same protein target) as a probabilistic model without using any machine learning model, called the *average interaction* method. Additional imputations and randomization tests were applied and compared to better analyze the basic concept of SIMPA (see **Supplementary Note 7**).

Because of the better resolution, available for H3K4me3 processed as genomic bins of size 5 kb, we present below results on this histone mark and refer to supplementary material for H3K27me3 processed at 50 kb bins (**Fig. S6**). For benchmarking, we used a single-cell ATAC-seq imputation method, SCALE, solely based on the single-cell dataset itself (*reference-free*) in contrast to SIMPA, which takes advantage of information from the reference bulk dataset. After applying a two-dimensional projection on the sparse and imputed datasets, we observed that the separation between the cell types was retained by SIMPA and by the reference-free method, contrary to the average interaction method (**Fig. 5A**). Moreover, three T-cell outliers were successfully associated to the related cell-type cluster by SIMPA, which achieved a slightly better homogeneity of the clusters in comparison to SCALE (**Fig. S6**). Dimensionality reduction was done by a combination of principal component analysis (PCA) and t-stochastic neighbor embedding (t-SNE) as suggested by Grosselin *et al*. on their analysis of sparse data (Grosselin *et al*., 2019). Different to the suggested procedure, we excluded the cell filtering, as we were interested to observe outliers after imputation.

**Figure 5.**
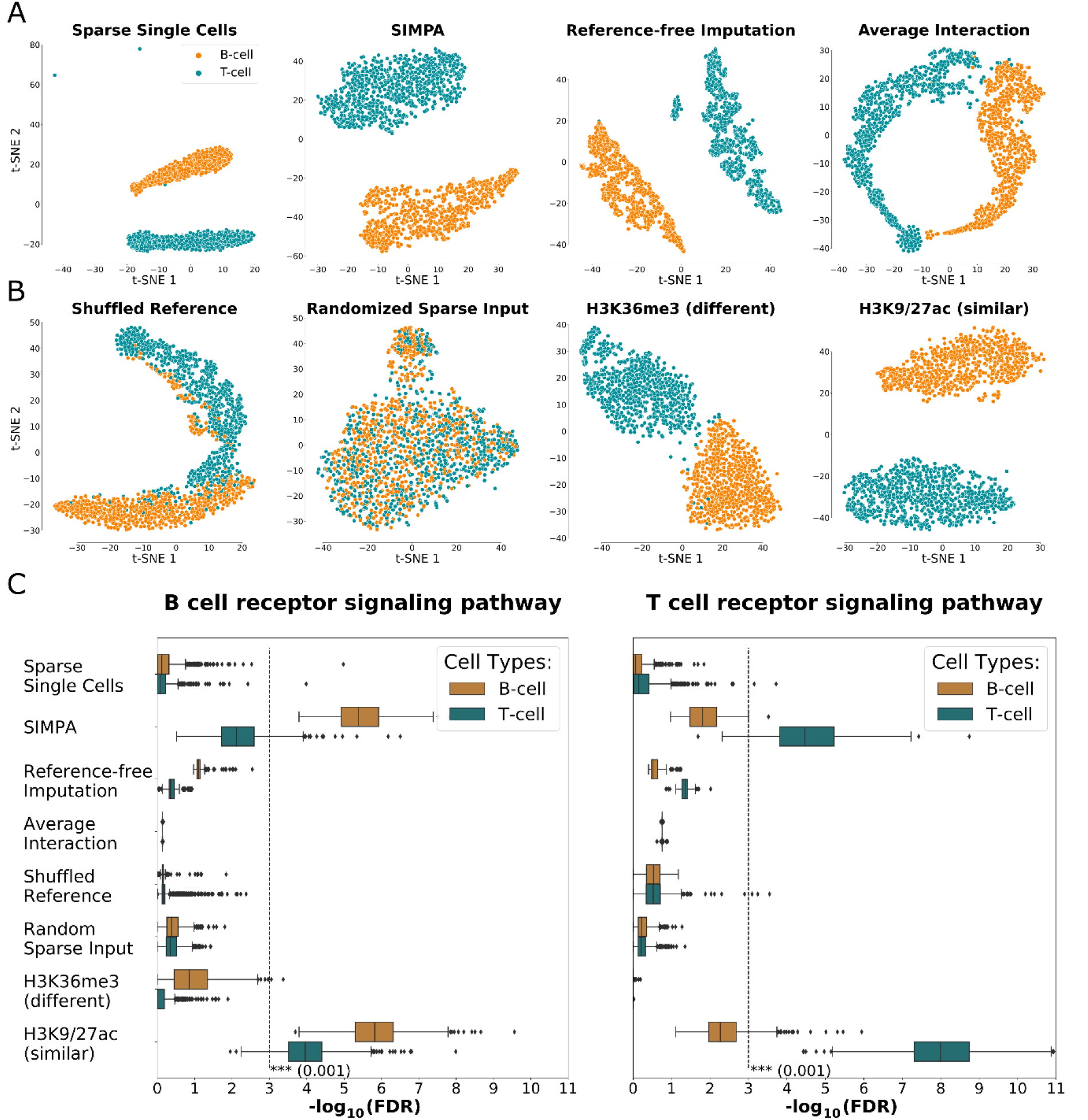
Cell-type specificity validation. **A + B Separation of single cells according to cell type. A**. Dimensionality reduction analysis applied on the H3K4me3 data derived from (i) the sparse single-cell data and three different imputation methods, (ii) SIMPA, (iii) reference-free imputation, and (iv) average interaction based on expected frequencies in the reference set. Results from SIMPA and from the reference-free method achieve the best clustering by separating the single cells (points) by cell types (colors). **B**. Effects of input modification on SIMPA, (i) using a shuffled reference set or (ii) randomized sparse input data, or using other histone marks as reference instead of H3K4me3, either (iii) the functionally different histone mark H3K36me3, or (iv) the functionally similar histone marks H3K9ac and H3K27ac. **C. Pathway enrichment analysis**. Boxplots show the significance of pathway enrichment analyses of genes annotated by single-cell regions as log-transformed false discovery rate (FDR; x-axis). Each dot represents the FDR of one single cell from the results of the different analysis experiments shown in **A+B** (y-axis). The dashed lines represent the log-transformed significance threshold of an FDR equal to 0.001. Only SIMPA achieves significant results by imputing preferably genomic regions associated with relevant pathway-related genes.

In order to validate further the algorithmic concept of SIMPA, we implemented two randomization tests in which either the ENCODE reference information was shuffled (Shuffled Reference) or the sparse single-cell input was randomly sampled (Randomized Sparse Input). Additionally, we applied SIMPA on the same data but with different histone marks as target. The selected histone marks were H3K36me3, a histone mark functionally different to H3K4me3, and H3K9ac and H3K27ac, a group of two histone marks functionally related to H3K4me3. These two marks were used together to increase the training data size. From this comparison, we observed that (i) the separation on the projection is lost after removing statistical patterns through shuffling or randomization, (ii) separation quality is moderate with an input mark functionally different from the real mark, and (iii) separation quality stays high using SIMPA with target histone marks functionally similar to the real mark (**Fig. 5B**). Thus, the most relevant statistical patterns from the reference dataset are identified by both the selection of single-cell-specific regions and the selection of target-specific experiments. Similar observations were made for H3K27me3 although a more compact clustering could be achieved on the SIMPA profiles compared to those from the reference-free method (**Fig. S6**).

Across several dimensionality reduction procedures applied for the H3K4me3 dataset, SIMPA and the reference-free method were both stable in retaining the cell-type clustering (**Fig. S7**). From these analyses we additionally conclude that the UMAP method using the Jaccard-Index distance achieves reasonable results when applied directly on the sparse data (in comparison to the common approach in single-cell analysis that uses first a PCA to select dimensions).

As pathway enrichment analysis is a common step in ChIP-seq data exploration, we next investigated if enrichment analyses of cell-type-specific pathways for individual single cells improve after applying imputation. We analyzed the sparse profiles and different imputed results with the KEGG pathway analysis function of the Cistrome-GO tool (S. Li *et al*., 2019). As reported in (**Fig. 5C**), the original sparse data did not provide enough interaction sites to show a significant pathway enrichment for any of the two cell types. Results from the reference-free strategy showed an improvement but not significant. However, with regions imputed by SIMPA, it was indeed possible to achieve significant enrichment scores and recover the cell-type-specific pathways for most of the cells. These results show that SIMPA is able to integrate functionally relevant information from the reference data in order to impute additional biologically meaningful regions, in contrast to the reference-free method, which is limited to the single-cell dataset.

### Optimal size of the imputation sets

As described, SIMPA computes the imputed probabilities for numerous genomic bins, and sorts and prioritizes those accordingly for imputation. In the previous validations, as a default, we imputed a number of bins equivalent to the average number of bins observed across all bulk ChIP-seq profiles from the target-specific reference set. On 5 kb resolution, the average number of bins of the H3K4me3 experiments is 32,584. However, once the bins are ranked by the imputed probability it is up to the user to alternatively create imputed sets of different sizes. With the next analysis we address the question about the optimal number of bins needed to improve the cell-type clustering at the same time enabling the detection of the relevant biological function by a significant enrichment of the correct pathway (for details see **Supplementary Note 8**).

The best cell-type clustering quality, evaluated by the Davies-Bouldin score (Davies and Bouldin, 1979), is reached when adding ∼11,000 bins (**Fig. 6A**). At this level, SIMPA slightly improves the clustering quality compared to the reference-free method.

**Figure 6.**
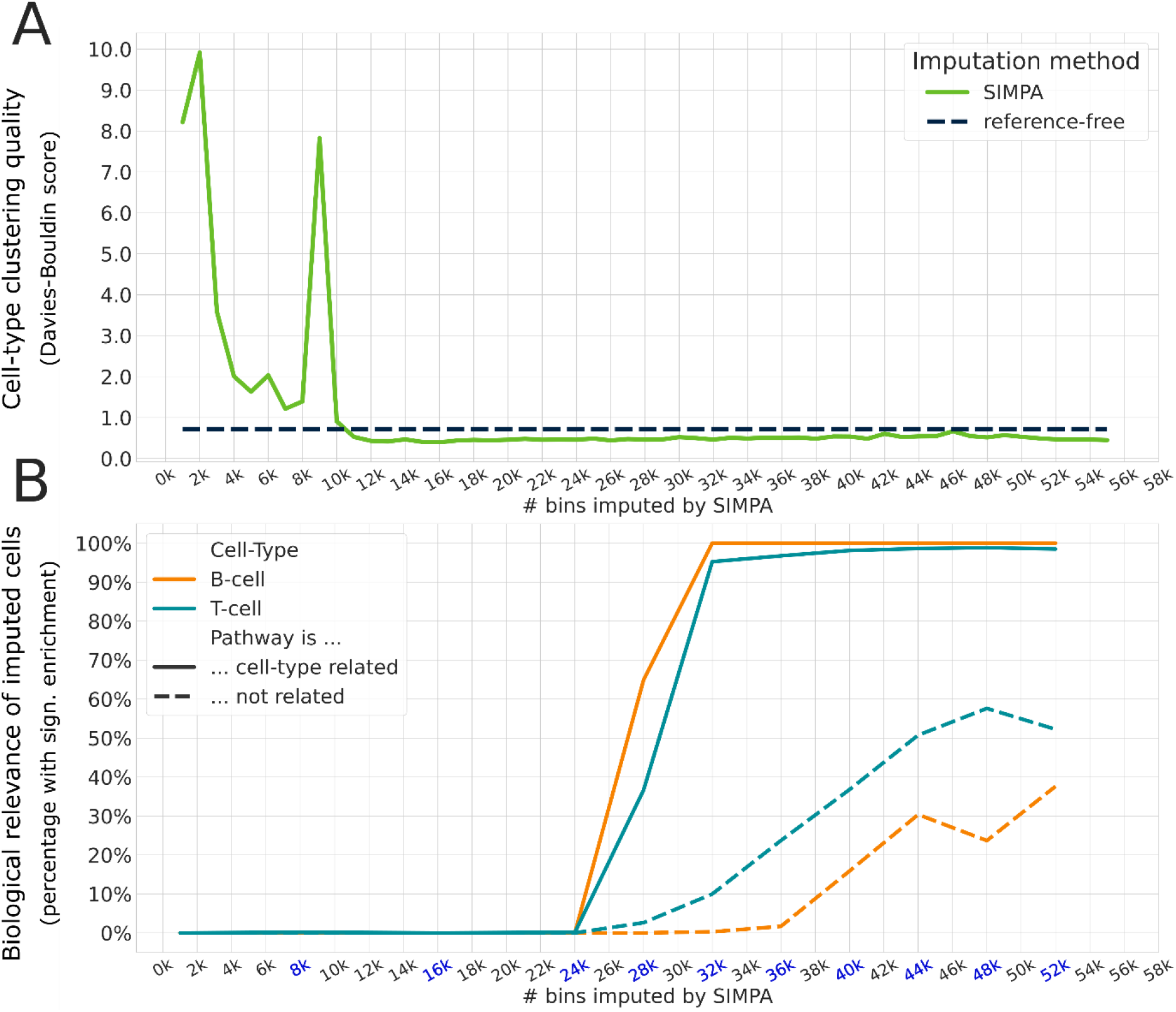
Clustering quality and pathway enrichment for different sizes. **A**. Clustering quality (y-axis) evaluated with the Davies-Bouldin score (the lower the better) applied on the imputed data after dimensionality reduction as described for Fig. 5 A+B. While the reference-free method derived only one imputed set for all the single cells (dashed black line), we could derive several imputed sets of different sizes using the imputed probabilities from SIMPA (x-axis). **B**. The pathways under investigation are the B-cell and T-cell receptor signaling pathways. In this way we analyze two pathways surely related or unrelated with the cell-types present in the dataset, B-cell and T-cell. The y-axis describes the percentage of imputed profiles for which a significant enrichment of the aforementioned pathways could be achieved. The dashed lines represent cases for which the unrelated pathway is significantly enriched, which is the T-cell receptor signaling pathway when analyzing a B-cell and vice versa. The significance level used is 0.001 similar to the analysis shown in Fig 5C. To reduce the computational resources spent on the pathway enrichment, this was done for ten imputation set sizes (highlighted in blue on the x-axis).

Considering the amount of cells in which the cell-type related pathway is significantly enriched, we observed that in ∼50% of the cells the related pathway is associated when adding ∼28,000 bins (**Fig. 6B**). After adding more than 32,000 bins, almost all cells have a significant enrichment for the cell-type related pathway, however, it seems to be also the limit for avoiding the association of the unrelated pathway. For this analysis the same pathways and settings are used as in Fig. 5C; the unrelated pathway is the T-cell receptor signaling pathway when analyzing a B-cell and vice versa.

## Discussion

After confirming the presence of statistical patterns within the ENCODE bulk ChIP-seq reference data, we show that machine learning models can leverage those patterns for the inference of interaction sites in sparse single-cell ChIP-seq profiles from individual single cells.

Based on the simulations, we could show this also for several cell-type-target combinations even if the experiments related to the cell-type were completely excluded from the training set. In both types of validation (leave-out origin and leave-out cell-type validations), SIMPA was able to capture cell-type-specific patterns even though the reference set was composed of profiles from many different cell-types and tissues, or the cell-type related data was completely excluded. Because the number of available bulk experimental profiles (ENCODE datasets) differs between targets, different training set sizes are available for different targets, with the smallest training set for H3K9ac (49 biosamples). Even for training sets of smaller size, the predictive performance remained high, although we expect models to be more reliable the larger the training set. Given that data portals such as ENCODE are still growing, we expect that the model reliability will increase in the future for many targets with a growing number of available reference datasets.

The interpretation of the SIMPA models, done with InterSIMPA applied on a real scChIP-seq dataset, allows us to reveal additional information from the ChIP-seq profiles measured within individual cells regarding regions responsible for the imputation. Importantly, leveraging reference data allows us to impute regions that were not present in the single-cell dataset at all, in contrast to a reference-free strategy. Considering for example the promoter regions of the T-cell receptor signaling pathway genes *CTLA4* and *ICOS*, these promoter-regions are not detected in any of the cells from the Grosselin *et al*. dataset, however, both have a high imputed probability from SIMPA. Moreover, for both promoters a high correlation coefficient was achieved within the validation by STRING co-expression values, confirming that our implementation not only answers the question about whether these promoters should be imputed or not, but it additionally reveals valuable information about regulatory relations implied by the single-cell dataset.

Regarding the full data imputation analysis, we observed further advantages of SIMPA’s reference-based imputation strategy compared to the reference-free imputation method. While both algorithms achieve a good separation of the cell types, only with the imputed profiles from SIMPA it was possible to determine the relevant biological function of single cells as shown by the pathway enrichment analysis. This suggests that SIMPA imputes biologically meaningful regions which are of functional relevance and confirms that, even though the training set involves a variety of different tissues and cell-types, SIMPA can find statistical patterns that belong to the correct cell-type. For single-cell datasets which reveal unknown subpopulations of cells, SIMPA could be used to identify active pathways for those cells after imputation. Interestingly, the quality of those results was maintained to some extent when not exactly the same scChIP-seq histone-mark target but functionally related targets were used to define the reference set. This suggests a valuable strategy to be applied for targets with a low availability of public bulk reference profiles.

SIMPA integrates solely datasets from bulk ChIP-seq in order to build the reference set. However, in the future, it will be relevant to integrate other types of data in order to complementarily extend the reference set. Such kinds of potential extensions are already in use.

For example, SCRAT is an analysis tool that summarizes single-cell regulome data using different types of public datasets such as genome annotations or motif databases that could be of interest for the application of SIMPA to transcription factor scChIP-seq profiles (Ji, Zhou and Ji, 2017). The scATAC-seq analysis tool SCATE performs imputation of missing regions integrating different types of public datasets (e.g. co-activated cis-regulatory elements and bulk DNase-seq profiles) (Ji *et al*., 2020). For future work, such approaches suggest the development of a reference-based method, allowing the imputation for both scChIP-seq and scATAC-seq data, integrating both types of reference data from the corresponding bulk assays and further complementary datasets.

SIMPA’s strategy, to train a model for each candidate bin and each single cell, results in its capability to produce highly relevant results and at the same time in its main limitation which is the requirement of a large amount of computational resources. Using a high-performance cluster, the results presented in this manuscript could be obtained within 1-2 days. However, if computational resources are limited, SIMPA offers the opportunity to run the imputation for a selection of cells which, for instance, represent a certain cluster to be analyzed. As shown, cell clusters can be identified even on the sparse profiles using the appropriate method for dimensionality reduction. Importantly, InterSIMPA can also be applied for individual cells, providing interpretable results within seconds of runtime.

## Conclusion

The strategy of SIMPA leveraging bulk ChIP-seq datasets for single-cell sequencing data imputation, is able to complete specifically sparse scChIP-seq data of individual single cells. In comparison to the non-imputed data and a reference-free imputation method, SIMPA was better at recovering cell-type-specific pathways. Furthermore, the interpretability of the machine learning models trained for the imputation can be used to reveal biologically important information from a sparse single-cell dataset. Conclusively, we developed an ensemble of computational methods to extract more information from a sparse dataset and impute missing data to better handle data sparsity of scChIP-seq datasets.

## Data and Software Availability

The ENCODE data used by SIMPA is available on the GitHub page of the software in preprocessed format. Single-cell ChIP-seq data used for the validations was taken from GSE117309. The SIMPA and InterSIMPA software are available at: https://github.com/salbrec/SIMPA. For more information about implementation details and the runtime see **Supplementary Note 9**.

## Authors’ contributions

All authors conceived the study and designed the experiments. SA and TA implemented the algorithm and validation analyses. All authors analyzed the data and interpreted the results. JF and MA co-supervised the research. All authors wrote the article.

## Acknowledgements

We thank Pablo Mier for proofreading the manuscript. We also thank the colleagues that tested SIMPA with their local machines: Kristina Kastano, Sweta Talyan, Jonas-Ibn Salem, and Gregorio Alanis-Lobato. Parts of this research were conducted using the supercomputer MOGON and advisory services offered by Johannes Gutenberg University Mainz (hpc.uni-mainz.de), which is a member of the AHRP (Alliance for High-Performance Computing in Rhineland Palatinate, www.ahrp.info) and the Gauss Alliance e.V. The authors gratefully acknowledge the MOGON supercomputer team. SA and TA thank the International PhD Programme (IPP) of the Institute of Molecular Biology, Mainz, for financial support. We thank Susanne Gerber and Leszek Wojnowski for meaningful discussions.

