## Supplementary Material for "Interpretable machine learning models for single-cell ChIP-seq imputation"

#### Supplementary Note 1 - Datasets

##### Preparation of the reference data (bulk ChIP-seq datasets)

To create the reference set that is used by SIMPA we downloaded all ChIP-seq experiments from the ENCODE portal that comply with the following criteria: the status is released, the experiment is replicated (isogenic or anisogenic), no treatment to the biosample, without genetic modification, and the organism is *Homo Sapiens* (human). For all experiments, we downloaded fully preprocessed sets of protein-DNA interacting regions as peak files: the *replicated peaks* for histone mark ChIP and the *optimal IDR thresholded peaks* for transcription factor ChIP. If possible, the peak files were downloaded for both assemblies hg19 and hg38. If one of the two was missing, we used the UCSC LiftOver tool to convert. Finally, we used 2251 experiments from different protein targets (antibody targets within the ChIP) and biosamples (either tissue or an immortalized cell line). For downloading, preprocessing, and updating the datasets we used a semi-automatic, SQL-backend procedure that was already used in a previous project (Andreani *et al.*, 2020).

##### Data preprocessing

In order to limit computational complexity, all reference experiments were converted from ChIP-seq peak sets to genomic bin sets. We provide on github the reference data in bin sizes of 5 kb and 50 kb for hg38 (https://github.com/salbrec/simpa). Bin sets for the reference ChIP-seq experiment are in binary format to be more efficiently integrated by the main Python scripts. Given one reference experiment, a bin is said to be "present" if there is at least one ChIP-seq peak that overlaps this bin, "absent" otherwise.

##### Preprocessing of scChIP-seq data (Grosselin *et al.*, 2019)

We downloaded the count matrices for H3K4me3 and H3K27me3 available in GEO under accession number GSE117309 in 5 kb and 50 kb binning resolution, respectively. From the matrices, we derived bed files for every single cell excluding gender-specific chromosomes. SIMPA, InterSIMPA and other imputation methods were then applied on 25% of the single cells randomly sampled, 1520 bed files for H3K4me3 and 1128 bed files for H3K27me3.

#### Supplementary Note 2 - The SIMPA algorithm

SIMPA is an algorithm implemented in Python 3.7.3 for "**S**ingle-cell Ch**IP**-seq i**MP**ut**A**tion", which is applied to one single cell represented by a sparse set of scChIP-seq genomic regions (or peaks) provided by the user in bed format. Within the algorithm, the given single-cell bed file is converted into a set of bins SC describing the single-cell input. The user also provides the target that is the name of the histone mark or transcription factor targeted by the antibody within the single-cell immunoprecipitation. The target is needed to specify the training set, which consists of experiments from the ENCODE reference set.

Unlike other single-cell imputation methods, SIMPA does not use information from other single cells. The imputation strategy is to use the sparse input from one given single cell to impute missing bins based on predictive information within bulk data between genomic regions bound by the target. In order to make the bulk data informative, first SIMPA collects all the ENCODE reference experiments available for the given target that defines the rows of the reference set matrix (*RS*) where columns represent bins:

$$RS = (a_{i,j}), 1 \leq i \leq n, 1 \leq j \leq m$$

*with*

$a_{i,j} \in \{0,1\}$ describing a cell of the matrix with value = 1 when bin $j$ in reference experiment $i$ is present, 0 otherwise,

and where $n$ is the number of experiments available for the given target, and $m$ is the number of bins that are present in at least one of the target specific experiments. As the rows are defined by the given target, the target-specificity is induced within this step.

Second, a subset of $RS$ is created by selecting only the columns for bins that are present in $SC$ to create the training features *TF*:

$$TF \subset RS,$$

$$TF = (a_{i,k}), 1 \leq i \leq n, 1 \leq k \leq s,$$

where $k$ indexes a selection of bins from $RS$ that are present in $SC$ and with $s$ the number of bins in $SC$ (see **Fig. 1B**). Bins present in $RS$ but not in $SC$ are collected and named as candidate bins $c$ that are potentially imputed bins.

Third, SIMPA takes each candidate bin in $c$ separately to compute an individual imputed probability $\rho_i$ for each $c_i$. Given $c_i$, SIMPA trains a classification model $cm_i$ based on $TF$ defining the features and $c_i$ as the class vector. Because an individual model is trained for each individual genomic bin, bin-specificity is induced for the whole approach. The imputed probability $\rho_i$ is finally computed by $cm_i$ which takes as input an artificial instance vector $a = (a_k), a_k = 1, 1 \leq k \leq s$. Consequently, $\rho_i$ is the probability of $c_i$ to be predicted for the imputed single-cell result, given the fact that all bins in $SC$ are observed. As we use a Random Forest implementation from the scikit-learn (version 0.21.3) Python's library (Pedregosa *et al.*, 2011; Buitinck *et al.*, 2013) with default settings to build classification models, the imputed probability is then the mean predicted class probability of the trees (by default 100) in the forest while the class probability of a single tree is calculated by the fraction of samples of the same class in a leaf.

Finally, SIMPA creates two files: one file in bed format and the other in SIMPA format described as a table listing the single-cell bins first, followed by the imputed bins sorted by the imputed probability. A line represents a bin described by its ID, its genomic coordinates, its frequency according to the target-specific reference experiments, and the imputed probability. Note that the first bins on top of this file have no imputed probabilities as they represent the original sparse single-cell input (a default value of -1 is assigned). The second file created by SIMPA is the imputed bed file containing the original single-cell bins and the additional imputed bins selected among those with the highest imputed probability. The number of bins within this bed file is defined by the average number of bins present in the target-specific bulk experiments, e.g. 32,584 for H3K4me3 (5 kb bin size) and 12,598 for H3K27me3 (50 kb bin size).

### Supplementary Note 3 - Cross validations

In order to validate whether machine learning models can be trained to accurately predict the observation of a bin, we applied the following approach: given the target, 10 single cells were randomly sampled for both cell types (B-cell and T-cell); for each single cell the training feature matrix $TF$ was created as explained for SIMPA while collecting also the candidate bins $c$. Then, for each candidate bin that defines the class vector, a Random Forest classification model was trained and evaluated by the area under the ROC-curve within a ten-fold cross-validation. In addition, we used the area under precision-recall curve to better study the class vector imbalance.

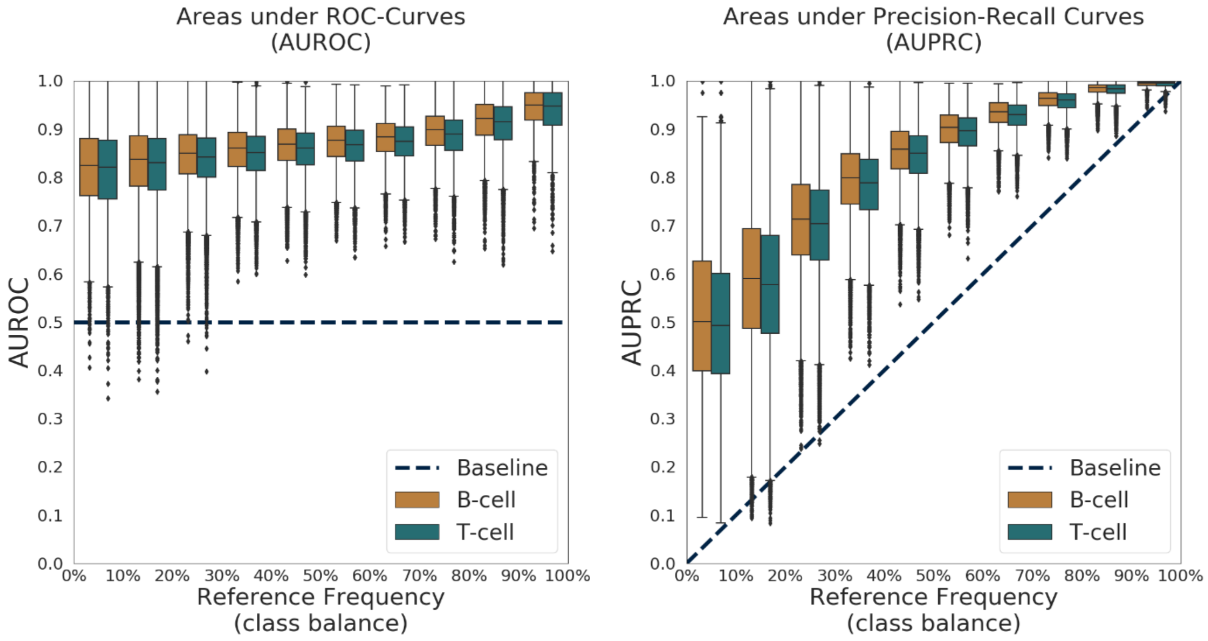


*Figure S1 - Cross-validations within H3K27me3 data*

*Cross validations within H3K27me3 data consisted of evaluating SIMPA's Random Forest performance to predict the values of each candidate bin of the single cells. For each bin, a ten-fold cross-validation was applied and summarized as Area under ROC-Curve (AUROC) or Area under Precision-Recall Curve (AUPRC) (y-axes). Results for all bins are presented by boxplots subdivided by class balance in the candidate bins (percentage of "1" values in the bin) (x-axis). The dashed lines describe the baseline performance expected from a random classification model: 0.5 for AUROC and equal to the class balance for AUPRC.*

#### Supplementary Note 4 - Recall of single-cell-specific bins

The goal of this analysis was to evaluate if SIMPA can recapitulate bins that are specific to individual single cells. It was important to analyze this, because the genomic regions detected for the reference experiments could be different to those detected by the single-cell approach. Given the H3K4me3 dataset, SIMPA was able to impute the vast majority of bins that were removed before imputation when using imputed sets of size ~32,000 which is similar to the average size of a bulk H3K4me3 experiment.

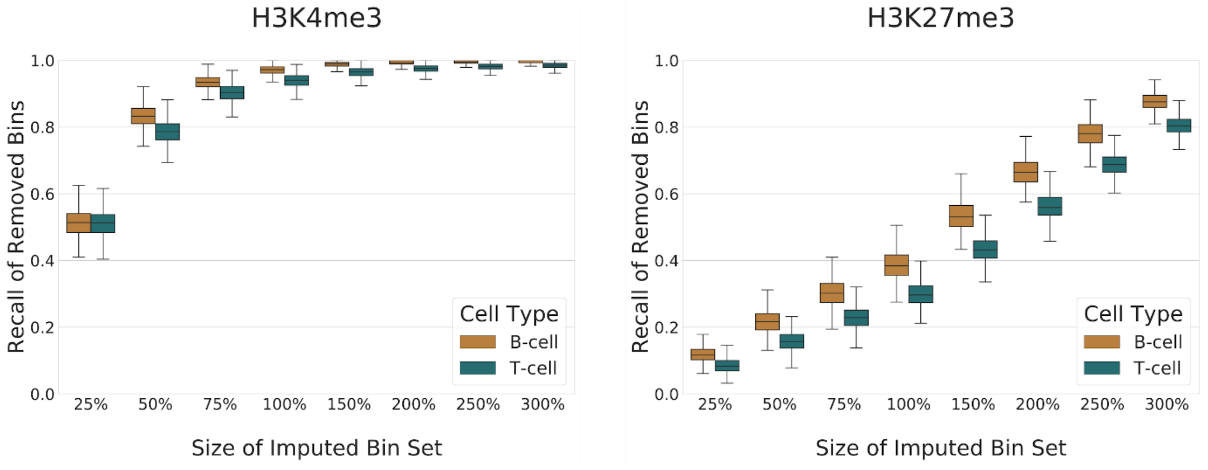


*Figure S2 - Recall of single-cell-specific bins*

*For every single cell of the H3K4me3 and H3K27me3 data, one third of randomly selected ChIP-seq genomic regions (or genomic bins) have been removed (set A) and the remaining two thirds used as input for SIMPA (set B). From the output of SIMPA, we calculated the recall of bins from set A (number of imputed bins from set A / total number of bins in set A). The boxplots summarize results for all single cells by recall values (y-axis) and the number of imputed bins as the percentage of its recommended size (x-axis). Considering the H3K4me3 data, approximately 90% of single-cell-specific removed bins could be recalled with the recommended size (100%).*

### Supplementary Note 5 - Simulated sparse interaction profiles

To apply and validate the imputation on several targets and different cell-types, sparse interaction profiles were simulated from the bulk experiments used by SIMPA as reference data. Using one bulk experiment as origin for the simulation, 11 sparse sets were created by randomly sampling genomic bins on 5 kb resolution. The size of these sparse sets range from 100 to 1600 in steps of 150. Minimum and maximum size for the simulations were derived from the size distribution of H3K4me3 single-cell profiles on 5 kb resolution (**Fig. S3A**). Initially, we expected a main bias to be present in the single-cell dataset where bins observed within the single-cell dataset would be driven by the overall reference frequency for a given target. The expectation was that bins that are more frequently observed within the reference data for different cell-types and tissues would be overrepresented in the single-cell profiles. In order to analyze the existence of this bias, we plotted the distribution of bin-frequencies for different bin sets defined by the accumulation of all single-cell profiles, pseudo bulk profiles (accumulating only those single-cell profiles for a given cell-type) for B-cell and T-cell and the corresponding bulk experiments from ENCODE (**Fig. S3B**). Compared to a true bulk experiment, the distributions of the reference-frequencies derived from the single-cell dataset(s) are similar. This indicates that we can exclude the aforementioned bias and another more sophisticated randomization for the simulation is not needed from this perspective.

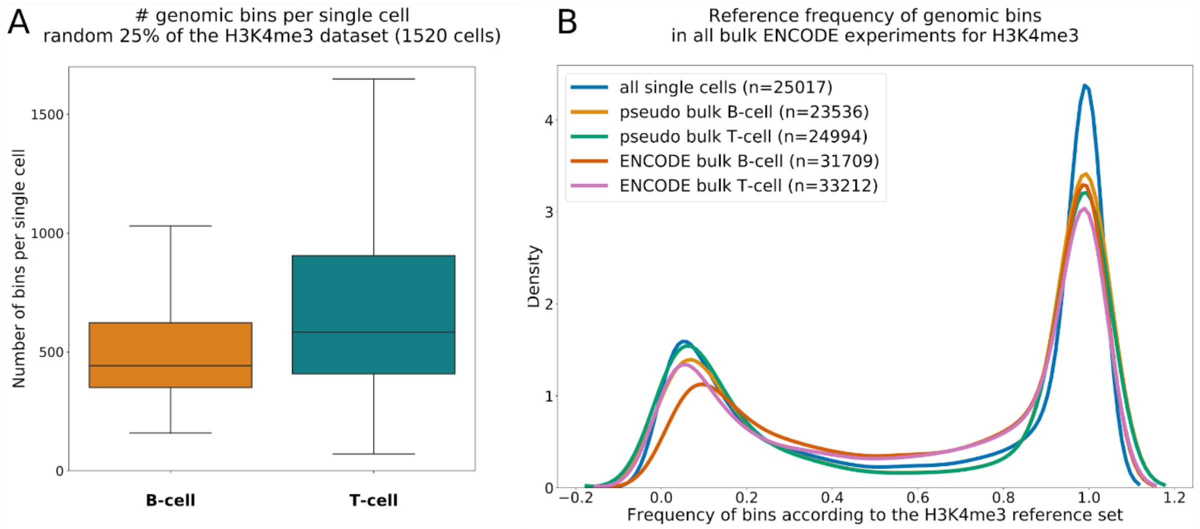


*Figure S3 - Observations based on the single-cell dataset for H3K4me3*

***A.** Boxplots showing the distribution of the number of bins within the sparse single-cell profiles separated by cell-type. **B.** Density of reference-frequencies derived from the H3K4me3 specific bulk ChIP-seq experiments for different bin sets: (i) all single-cell profiles accumulated, (ii) accumulated single-cell profiles for the two different cell-types, and (iii) profiles from ENCODE bulk ChIP-seq for the two cell-types. All bin sets contain numerous bins that are rather frequent according to the bulk ChIP-seq data (peak on the right-hand side). The peak on the left-hand side represents bins that are less frequently observed and might be more specific to the given cell-types.*

For each simulated sparse set, SIMPA was applied to calculate the imputed probabilities for the candidate bins. These probabilities were then used to plot the area-under ROC-curves (AUROC) in **Fig. 2 and S4** and area under precision-recall curves (AUPRC) in **Fig. S5**. The set of true interaction sites was defined by the interactions observed for the bulk ChIP-seq profile that was used as origin for the corresponding sparse simulated profile. By default, the origin was excluded from the training set used by SIMPA to train the classification models in order to perform a default validation, called *leave-out origin* (LOO). Another extreme validation was applied in which all reference experiments for the same cell-type were excluded, called *leave-out cell-type* (LOCT). For the AUPRC validation we added the random baseline defined by the fraction of positives in the origin bulk experiment used for validation. A more detailed description of the average interaction approach is given in **Supplementary Note 6**.

#### Analysis involving the consensus

As described in the main manuscript, one analysis involved the *consensus* profile of replicate experiments related to a same biosample-target combination. . For each group of at least 3 replicate experiments in the ENCODE reference data (e.g., defined by HepG2-CTCF combination), a simulation was ran for each replicate experiment taken as origin. The remaining replicate experiments were used to create the consensus profile by randomly sampling bins from the union of their bins. In order to exclude a bias from the imputed set sizes for the Jaccard-Index computation, we sampled the same amount of bins as present in the origin. In order to consider existing knowledge about the reliability of the bins within the remaining replicate experiments, the sampling was carried out with weights that ensure that bins present in multiple of the remaining experiments are picked with a proportional higher probability (e.g., 2 times more probable for bins present in 2 replicates compared to bins present in only 1 replicate, and 3 times more for 3 replicates). Before computing the Jaccard-Index, the bins from the simulated sparse profile were removed from all the sets, origin, imputed, and consensus. To avoid an imputed result being biased towards the origin or to the consensus, we ran this analysis only with the leave-out cell-type validation.

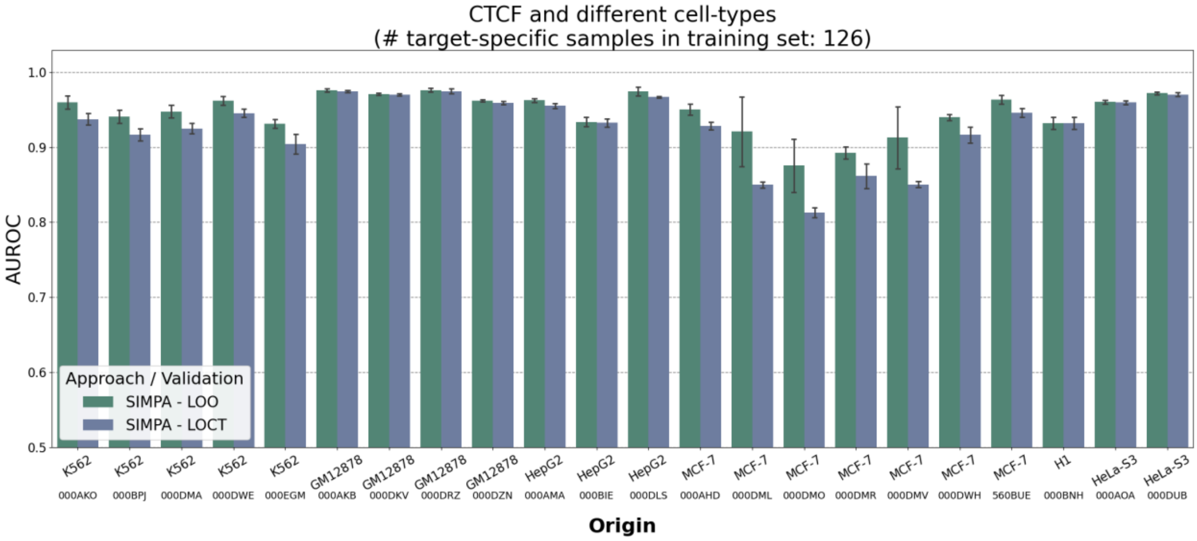
CTCF and different cell-types
(# target-specific samples in training set: 126)
AUROC
1.0
0.9
0.8
0.7
0.6
0.5
Approach / Validation
SIMPA - LOO
SIMPA - LOCT
K562 000AKO
K562 000BPJ
K562 000DMA
K562 000DWE
K562 000EGM
GM12878 000AKB
GM12878 000DKV
GM12878 000DRZ
GM12878 000DZN
HepG2 000AMA
HepG2 000BIE
HepG2 000DLS
MCF-7 000AHD
MCF-7 000DML
MCF-7 000DMO
MCF-7 000DMR
MCF-7 000DMV
MCF-7 000DWH
MCF-7 560BUE
H1 000BNH
HeLa-S3 000AOA
HeLa-S3 000DUB
Origin

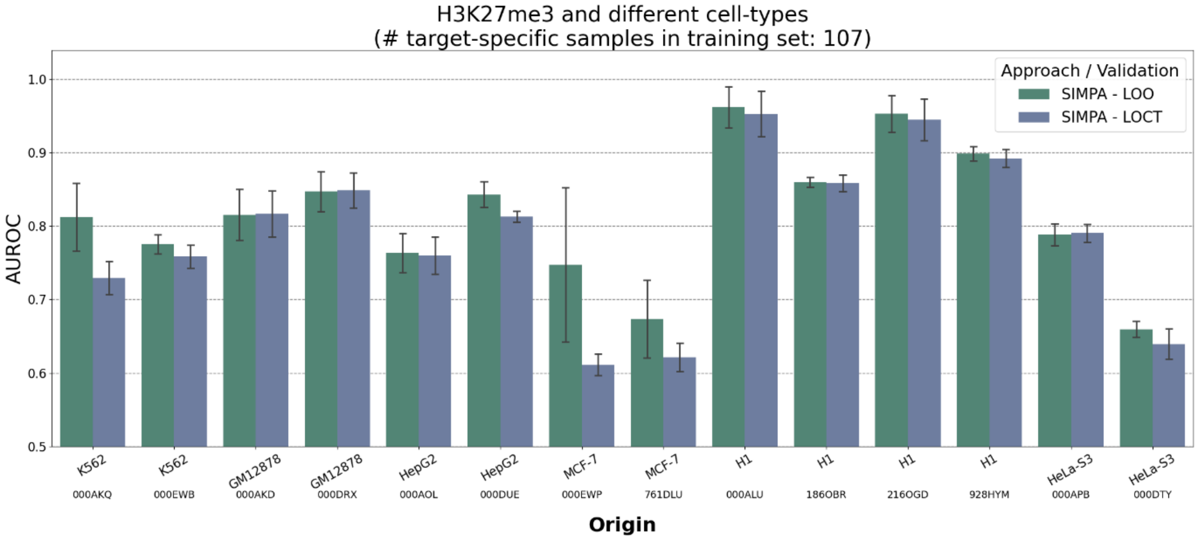
H3K27me3 and different cell-types
(# target-specific samples in training set: 107)
AUROC
1.0
0.9
0.8
0.7
0.6
0.5
Approach / Validation
SIMPA - LOO
SIMPA - LOCT
K562 000AKQ
K562 000EWB
GM12878 000AKD
GM12878 000DRX
HepG2 000AOL
HepG2 000DUE
MCF-7 000EWP
MCF-7 761DLU
H1 000ALU
H1 186OBR
H1 216OGD
H1 928HYM
HeLa-S3 000APB
HeLa-S3 000DTY
Origin

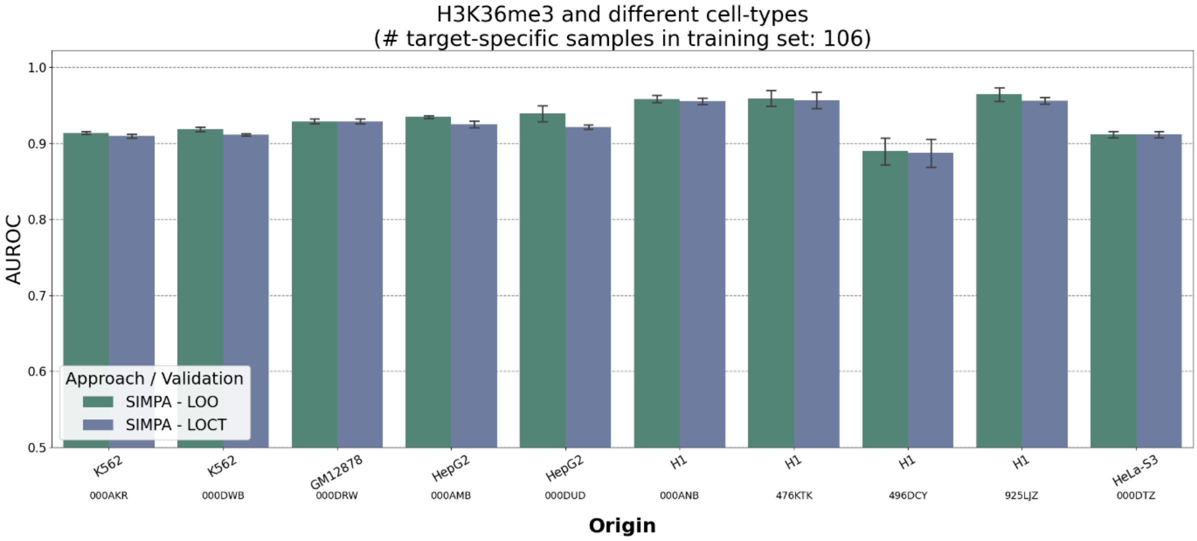
H3K36me3 and different cell-types
(# target-specific samples in training set: 106)
AUROC
1.0
0.9
0.8
0.7
0.6
0.5
Approach / Validation
SIMPA - LOO
SIMPA - LOCT
K562 000AKR
K562 000DWB
GM12878 000DRW
HepG2 000AMB
HepG2 000DUD
H1 000ANB
H1 476KTK
H1 496DCY
H1 925LJZ
HeLa-S3 000DTZ
Origin

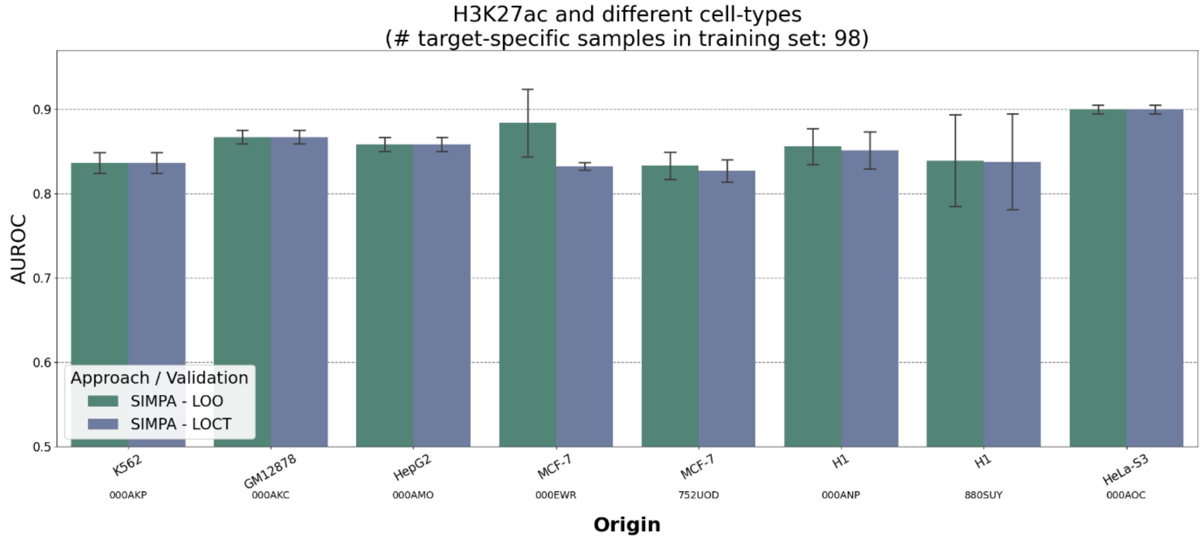
H3K27ac and different cell-types
(# target-specific samples in training set: 98)
AUROC
0.5
0.6
0.7
0.8
0.9
Approach / Validation
SIMPA - LOO
SIMPA - LOCT
K562
000AKP
GM12878
000AKC
HepG2
000AMO
MCF-7
000EWR
MCF-7
752UOD
H1
000ANP
H1
880SUY
HeLa-S3
000AOC
Origin

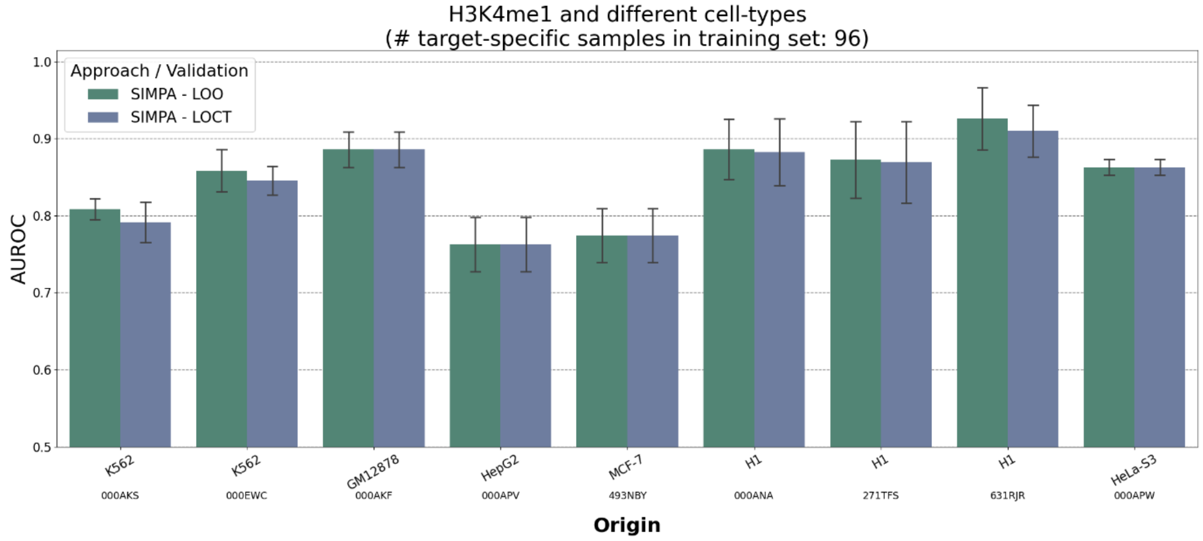
H3K4me1 and different cell-types
(# target-specific samples in training set: 96)
AUROC
0.5
0.6
0.7
0.8
0.9
1.0
Approach / Validation
SIMPA - LOO
SIMPA - LOCT
K562
000AKS
K562
000EWC
GM12878
000AKF
HepG2
000APV
MCF-7
493NBY
H1
000ANA
H1
271TFS
H1
631RJR
HeLa-S3
000APW
Origin

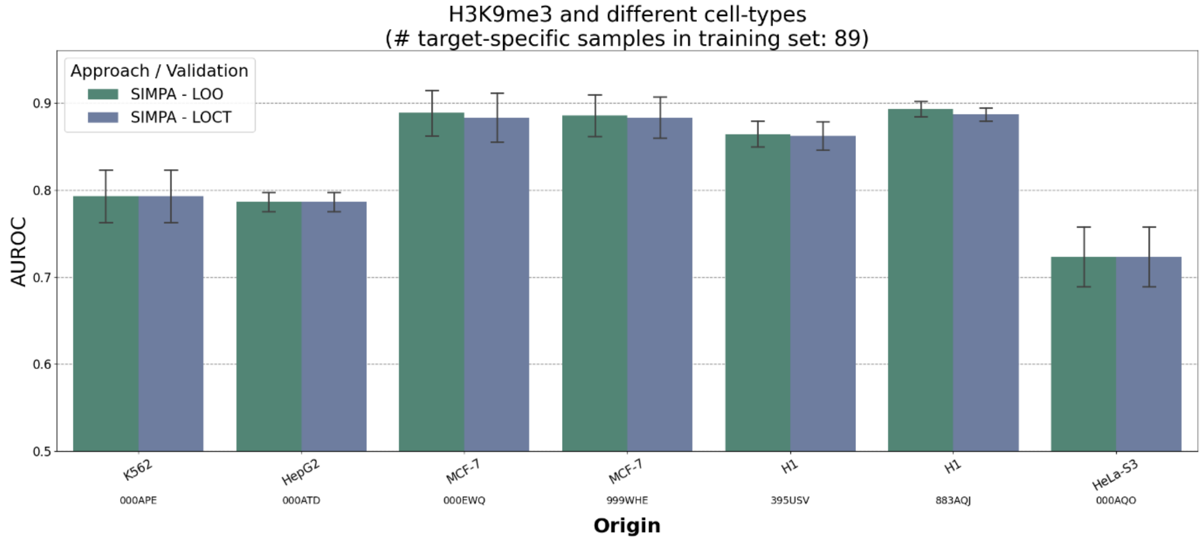
H3K9me3 and different cell-types
(# target-specific samples in training set: 89)
AUROC
0.5
0.6
0.7
0.8
0.9
Approach / Validation
SIMPA - LOO
SIMPA - LOCT
K562
000APE
HepG2
000ATD
MCF-7
000EWQ
MCF-7
999WHE
H1
395USV
H1
883AQJ
HeLa-S3
000AQO
Origin

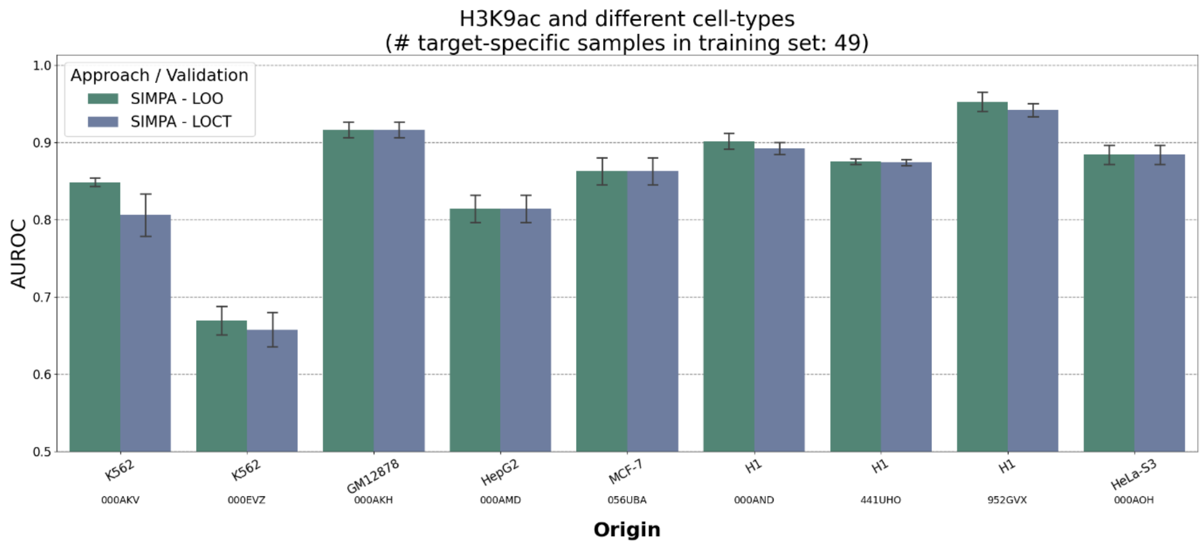


*Figure S4 - Area under ROC-curve (AUROC) on simulated sparse profiles (the remaining targets in addition to Fig. 2)*

*Using the origin for simulation as the validation set of true binding interactions, the area under ROC-curve (AUROC in y-axis) describes the capability of SIMPA to accurately impute and recapitulate the bulk experiment. The bars describe the mean AUROC achieved on multiple simulated profiles and the error bars describe the standard deviation across those multiple applications on sparse sets with different sizes. SIMPA was validated with two strategies, the default leave-out origin (LOO) and the extreme leave-out cell-type (LOCT). The x-axis labels indicate the cell-type and ENCODE accession of the origin profile. Note that, when only one bulk dataset for a specific target - cell-type combination is available in ENCODE (e.g., CTCF - H1) there is no difference between the two leave-out validations.*

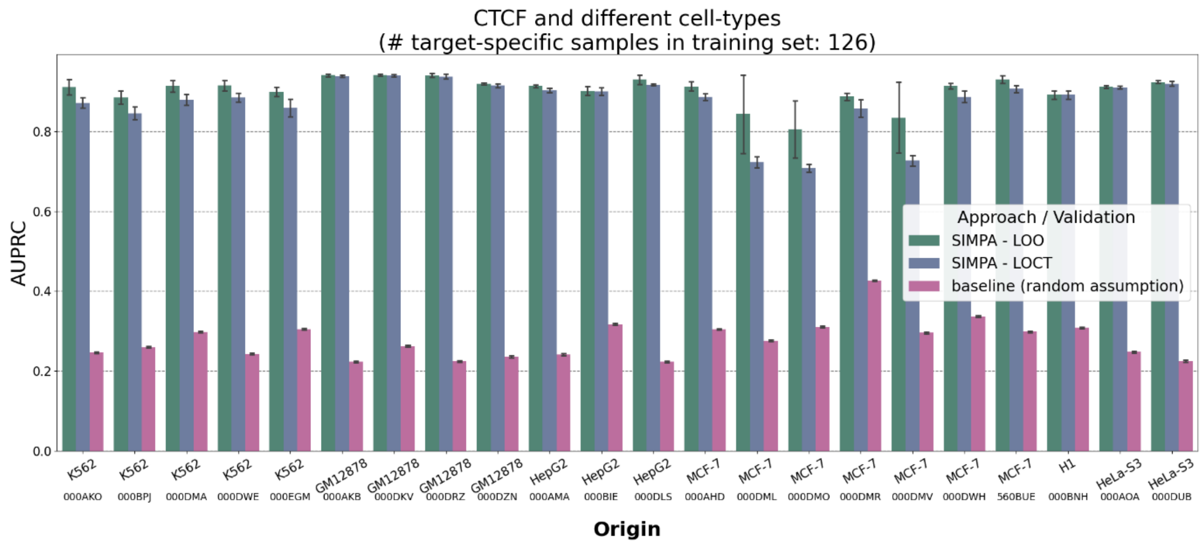

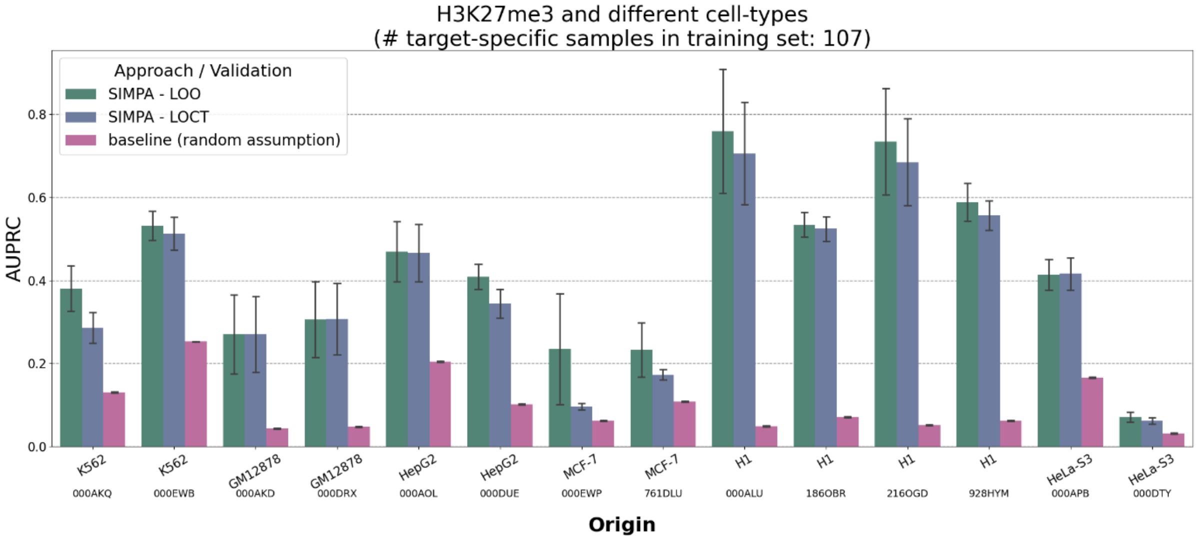
H3K27me3 and different cell-types
(# target-specific samples in training set: 107)
Approach / Validation
SIMPA - LOO
SIMPA - LOCT
baseline (random assumption)
AUPRC
0.0
0.2
0.4
0.6
0.8
K562 000AKQ
K562 000EWB
GM12878 000AKD
GM12878 000DRX
HepG2 000AOL
HepG2 000DUE
MCF-7 000EWP
MCF-7 761DLU
H1 000ALU
H1 186OBR
H1 216OGD
H1 928HYM
HeLa-S3 000APB
HeLa-S3 000DTY
Origin

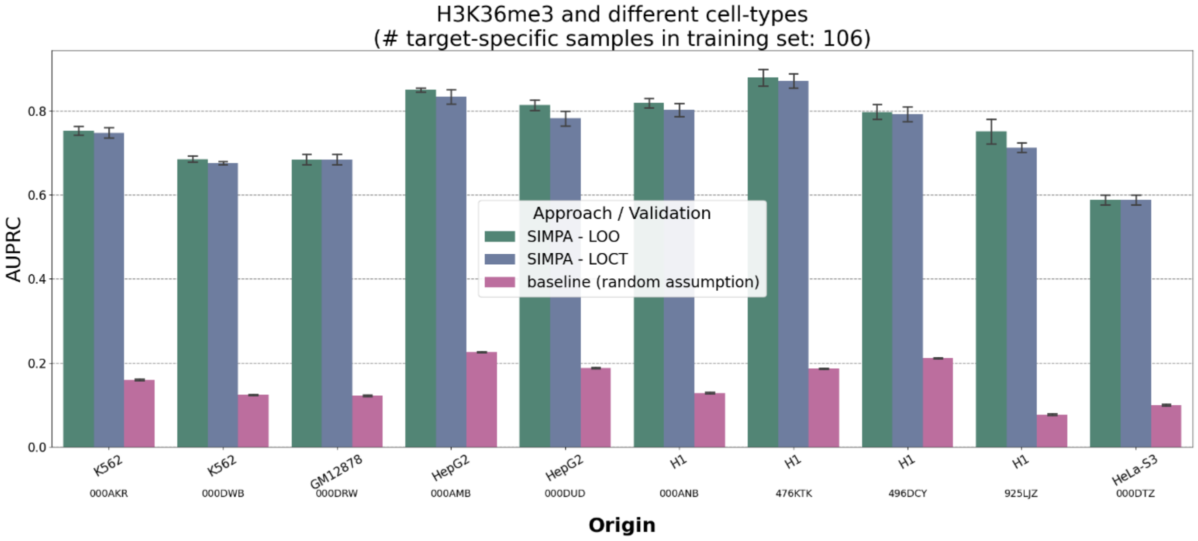
H3K36me3 and different cell-types
(# target-specific samples in training set: 106)
Approach / Validation
SIMPA - LOO
SIMPA - LOCT
baseline (random assumption)
AUPRC
0.0
0.2
0.4
0.6
0.8
K562 000AKR
K562 000DWB
GM12878 000DRW
HepG2 000AMB
HepG2 000DUD
H1 000ANB
H1 476KTK
H1 496DCY
H1 925LJZ
HeLa-S3 000DTZ
Origin

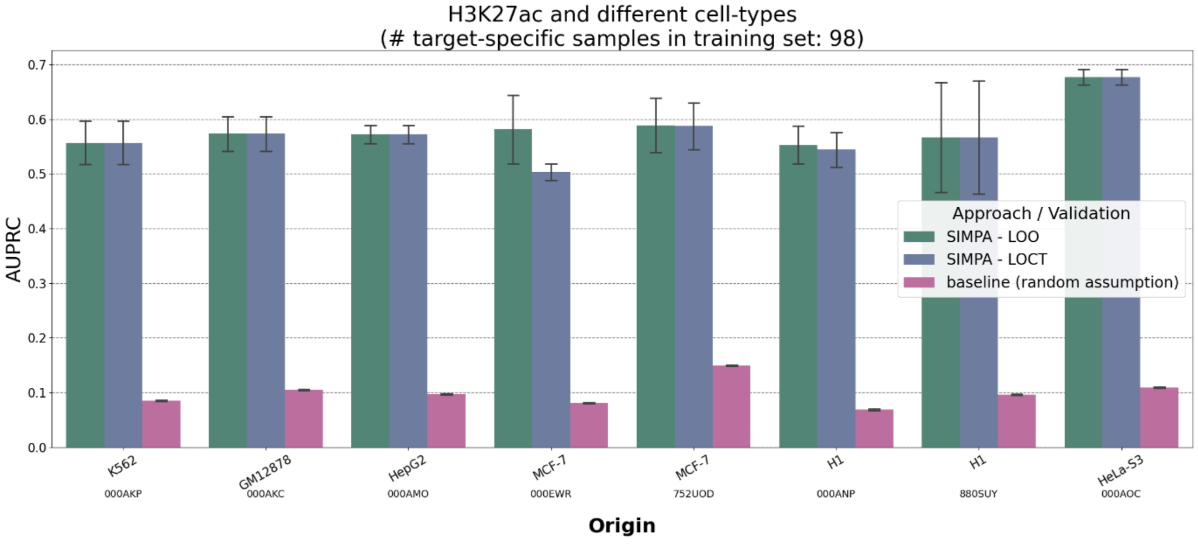
H3K27ac and different cell-types
(# target-specific samples in training set: 98)
Approach / Validation
SIMPA - LOO
SIMPA - LOCT
baseline (random assumption)
AUPRC
0.0
0.1
0.2
0.3
0.4
0.5
0.6
0.7
K562 000AKP
GM12878 000AKC
HepG2 000AMO
MCF-7 000EWR
MCF-7 752UOD
H1 000ANP
H1 880SUY
HeLa-S3 000AOC
Origin

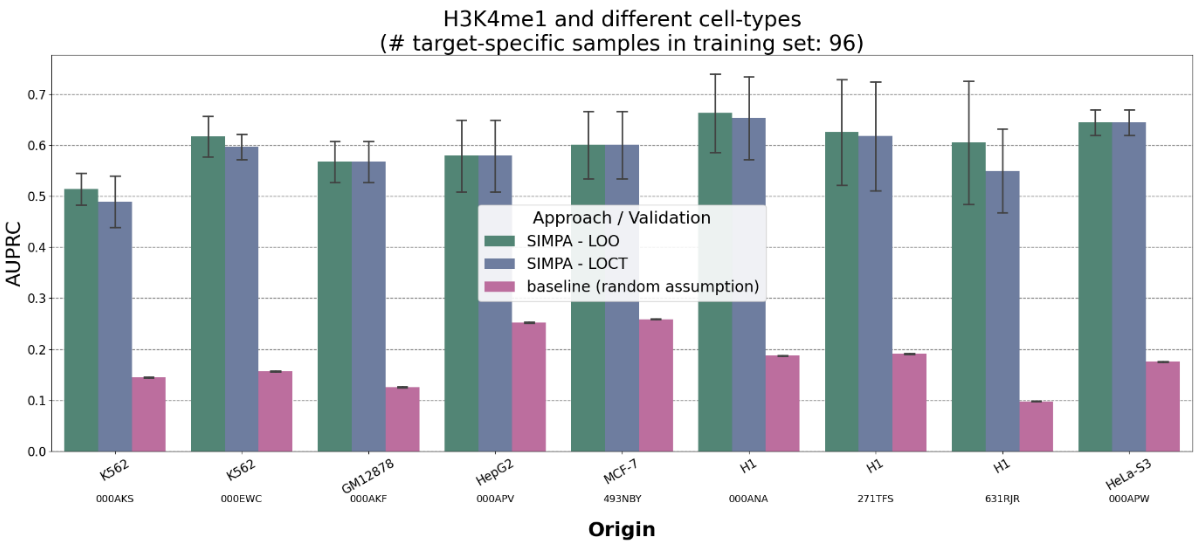


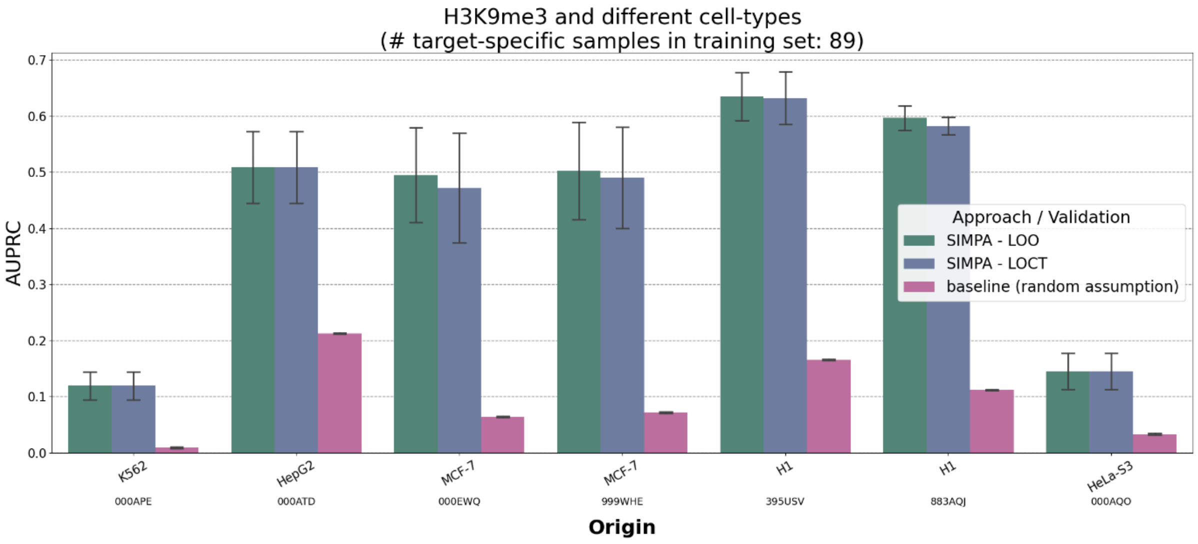


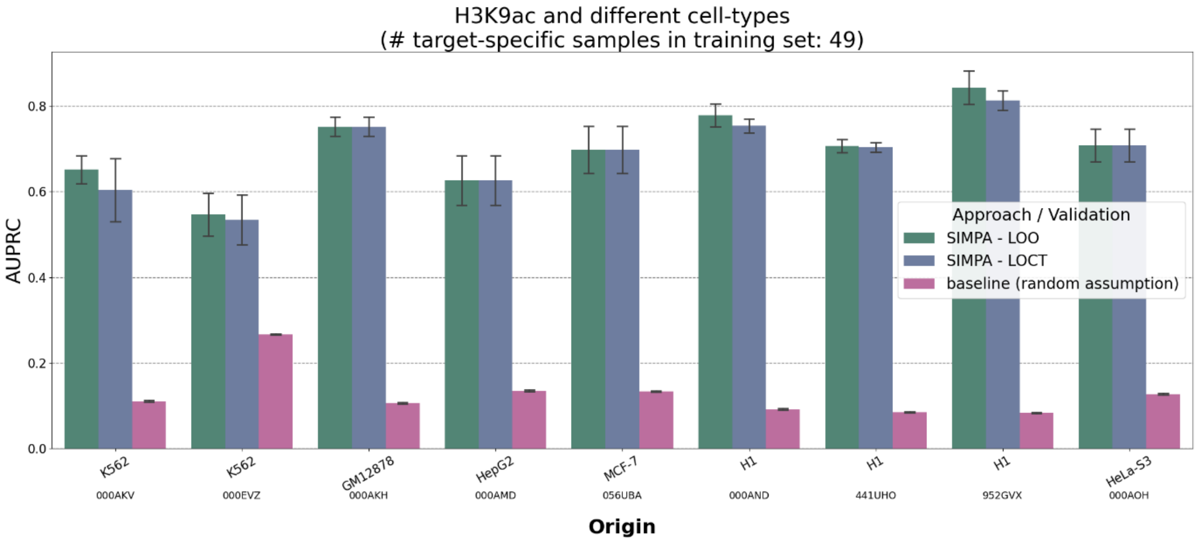


*Figure S5 1 - Area under precision-recall curve (AUPRC) on simulated sparse profiles (the remaining targets in addition to Fig. 2)*

*Same as in **Fig. S4** but using the area under precision-recall curve (AUPRC in y-axis) as performance measure. The pink bars show the class balance (fraction of positives in the class feature) representing the random assumption as baseline to be expected from a primitive classifier that randomly assigns the class values (according to* (Saito Takaya AND Rehmsmeier, 2015)*).*

### Supplementary Note 6 - Validation of model interpretation

#### The InterSIMPA algorithm

InterSIMPA applies the same imputation strategy as SIMPA but it trains only one machine learning model for one genomic region defined by the user via an additional mandatory parameter. Hence, given the single cell and the target, the training features are prepared in the very same way and there is also no difference with respect to the composition of the candidate bins. The additional parameter is a genomic position, defined by chromosome number and nucleotide position, which could represent the summit of a relevant ChIP-seq peak or simply the center of the region of interest for the user. This region of interest defines the candidate bin that contains the labels for the class used by the classification algorithm. The classification algorithm used is again Random Forest with default settings, but training 1000 trees. After training the model, the training features are ranked by the Random Forest feature importance. Training features are essentially genomic regions that are then displayed together with the closest gene, the distance to its TSS position, and some essential information about the gene from the RefSeq database (Pruitt *et al.*, 2014).

#### Interpretability analysis including the STRING database

To validate if the feature importance values returned by InterSIMPA have biological relevance, we applied InterSIMPA on the 1520 H3K4me3 single-cell profiles from B-cells and

T-cells from Grosselin *et al.* (explained above). For each single cell we applied InterSIMPA for several genomic positions representing the promoter regions of 120 selected genes. Single-cell promoter combinations for which an interaction was present were skipped for this analysis. From these genes, 67 and 97 are involved in the B-cell receptor and T-cell receptor signaling pathways, respectively. The two pathways have 44 genes in common. Each genomic position that was eventually used as input for InterSIMPA was defined by the center of the genomic window -1500 bp and +150 bp from the transcription start site (TSS) of the gene. As explained above, InterSIMPA returns the imputed probability for the given promoter position and the feature importance values of other observed regions from the given single-cell profile. Moreover, each region is automatically annotated by the closest gene. Given an InterSIMPA run for a specific promoter, these gene annotations were used to compute a value that describes the gene-gene relation with respect to the promoter's gene. This final value is the average feature importance across all single cells used within this analysis. Note, that usually we expect that InterSIMPA is used to annotate and report only regions with a high feature importance value according to the users' interest. However, for this analysis, all regions from the single-cell profiles were used without any restriction.

The STRING database provides fully preprocessed lists that include co-expression values for pairs of genes. Given one gene from the list of selected pathway-related genes, we first selected all other genes for which a STRING co-expression value and an InterSIMPA feature importance value were available. This results in two vectors. We used the Pearson's correlation coefficient between these two vectors to validate whether the predictive information of H3K4me3 interactions within the single cells reflect the co-expression of the annotated genes.

#### Supplementary Note 7 - Cell-type clustering and pathway enrichment analysis

##### Applied methods – dimension reduction and pathway analysis

In order to apply the dimension reduction, following the procedure from Grosselin *et al.*, Principal Component Analysis (PCA) was applied to compute the first 50 principal components. This first step was applied on a matrix that has one single cell in a row and the bins described by the columns, while bins are either solely from the sparse dataset or from imputed profiles. t-distributed stochastic neighbor embedding (t-SNE) was then applied on the first 50 principal components to reduce this matrix to two dimensions. We used PCA with default settings and t-SNE with 1000 iterations and a perplexity of 40. For both methods we used the implementations from the Python scikit-learn package. Different to the procedure from Groseelin *et al.,* we did not filter any of the single cells.

Pathway enrichment analysis on sparse and imputed bin sets was performed using the *cistromego* tool downloaded from https://github.com/changxinw/Cistrome-GO and applied with default settings. The Cistrome-GO results were then analyzed with focus on the B-cell receptor signaling pathway (KEGG: hsa04662) and the T-cell receptor signaling pathway (KEGG: hsa04660). We associated those pathways as the *cell-type related* pathways for the B-cells and T-cells, respectively. Considering Figure 5B of the main manuscript, the opposite pathway was used as *unrelated*.

##### Reference-free imputation

For reference-free imputation we used SCALE (a method developed for scATAC-Seq that uses the whole single-cell dataset for imputation; (Xiong, Lei, et al. 2019)) with default settings for the count matrices of H3K4me3 and H3K37me3 excluding gender-specific chromosomes and keeping all single cells. We used the flag --*binary* to obtain a binary description of the presence of single bins within the imputed results for each single cell.

#### Reference frequency and average interaction method

The reference frequency $freq$ of a particular bin $j$ describes its presence in the reference experiments in $RS$:

$$freq_j = \frac{\sum_{i=0}^{n} a_{i,j}}{n}$$

where $a_{i,j} \in RS$ describes the value of bin $j$ in experiment $i$, n being the total number of target-specific experiments in *RS*.

The reference frequency described above is generally a good indicator of the presence of a bin. Intuitively, the higher the frequency, the more likely the presence of the bin. We use this frequency to implement the baseline average interaction strategy. Hence, the average interaction strategy outputs the sparse bins from the single cell, plus imputed bins ranked by the reference frequency. By default, the number of bins for the imputed result is the average number of bins observed for the target-specific reference experiment.

#### Randomization tests

We randomized the reference set obtained from the mark of interest from the bulk ENCODE data maintaining the same frequency observed for each bin. For this operation, we used the numpy shuffle function in Python (numpy 1.17.2). The second randomization was performed by simulating the single-cell bins as a set of random sequences with an amount, length and nucleotide distribution similar to that observed for each single cell. For this operation we used the function “getNullseqs” from the package gkmSVM in R (Ghandi *et al.*, 2016).

#### Similar and different mark imputation

To analyze the impact of the selected target that defines the training set, we analyzed the imputed results using histone marks with either similar or different functionality than the actual histone mark. For H3K4me3, an activating mark, the selected similar marks were H3K9ac and H3K27ac with 49 and 98 available experiments, respectively, and the selected different mark was H3K36me3 with 106 available experiments. For H3K27me3, we used H3K36me3 as similar, and H3K9ac and H3K27ac together as different marks. Using a collection of two similar marks (H3K9ac and H3K27ac) allowed us to increase the number of reference experiments used for training. The size of the reference sets for marks H3K4me3 and H3K27me3 are 178 and 107, respectively.

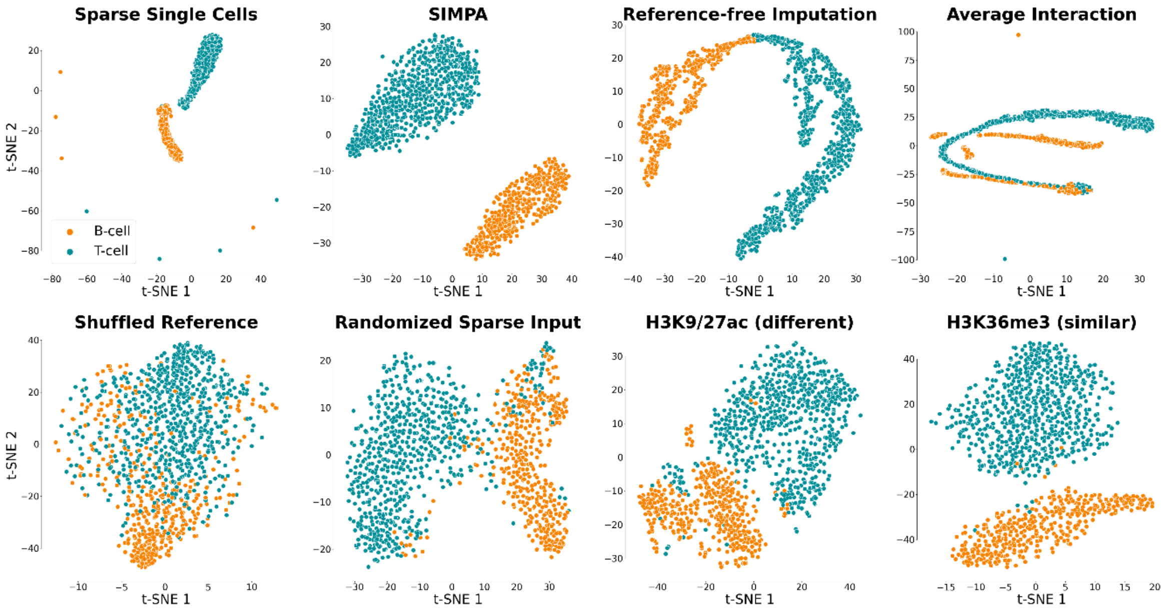


##### *Figure S6 - Cell-Type Specificity Validation of the H3K27me3 data*

*Separation of single cells according to cell type.* ***A*** *Dimension reduction analysis applied on the H3K27me3 data derived from (i) the sparse single-cell data and three different imputation methods, (ii) SIMPA, (iii) reference-free imputation, and (iv) average interaction based on expected frequencies in the reference set. SIMPA achieves the best imputation by maximizing the separation of single cells (points) by cell types (colors) without displaying the artifacts that are observed due to data sparsity (especially visible on the Sparse Single Cells plot).* ***B.*** *Effects of input modification on SIMPA, (i) using a shuffled reference set or (ii) randomized sparse input data, or using other histone marks as reference instead of H3K4me3, either (iii) the functionally different histone marks H3K9ac and H3K27ac, or (iv) the functionally similar histone mark H3K36me3.*

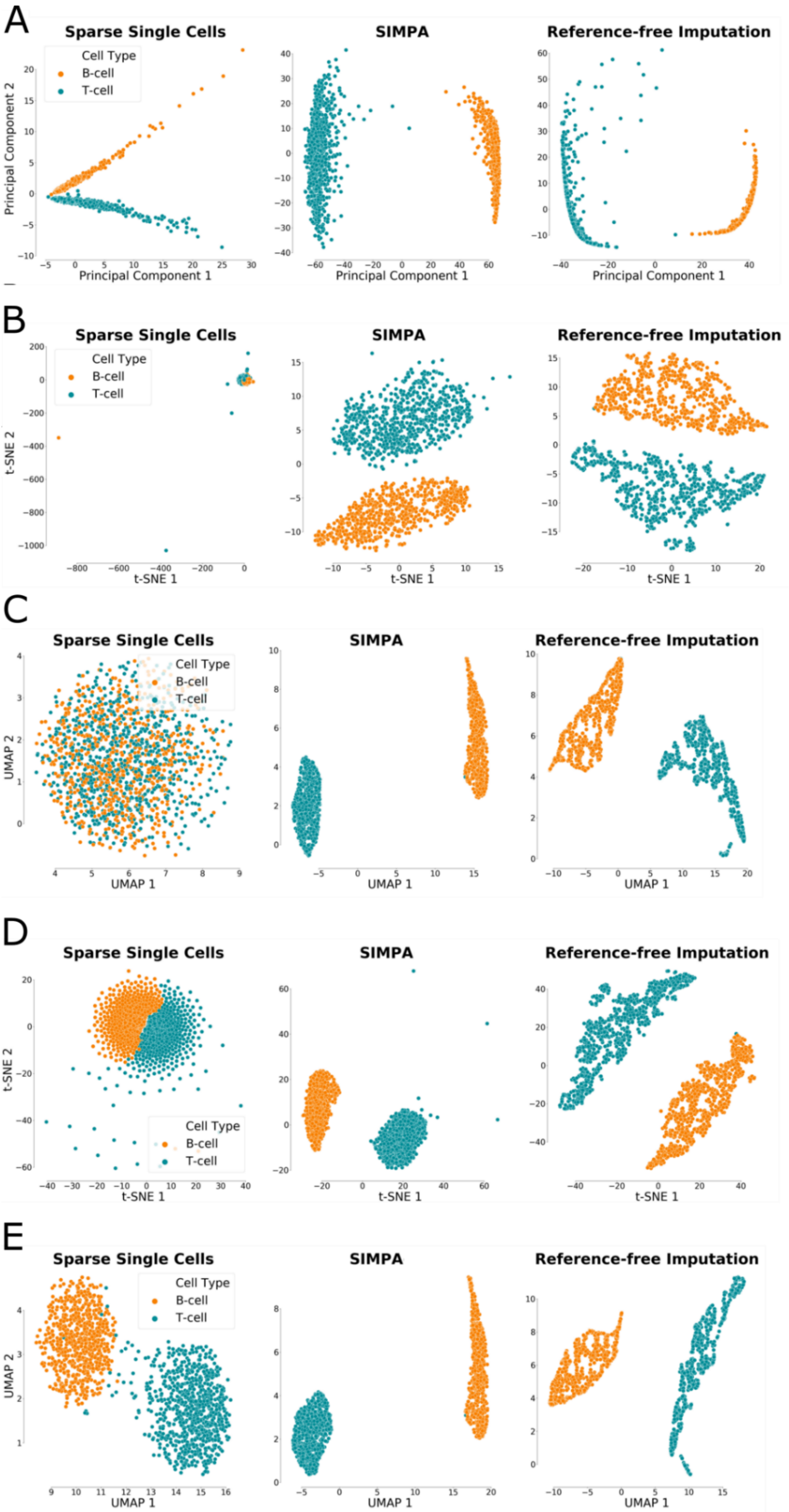

A
Sparse Single Cells
SIMPA
Reference-free Imputation
Cell Type
B-cell
T-cell
Principal Component 2
Principal Component 1
Principal Component 1
Principal Component 1
B
Sparse Single Cells
SIMPA
Reference-free Imputation
Cell Type
B-cell
T-cell
t-SNE 2
t-SNE 1
t-SNE 1
t-SNE 1
C
Sparse Single Cells
SIMPA
Reference-free Imputation
Cell Type
B-cell
T-cell
UMAP 2
UMAP 1
UMAP 1
UMAP 1
D
Sparse Single Cells
SIMPA
Reference-free Imputation
Cell Type
B-cell
T-cell
t-SNE 2
t-SNE 1
t-SNE 1
t-SNE 1
E
Sparse Single Cells
SIMPA
Reference-free Imputation
Cell Type
B-cell
T-cell
UMAP 2
UMAP 1
UMAP 1
UMAP 1

*Figure S7 - Several dimension reduction methods applied on the H3K4me3 profiles before and after imputation*

*Separation of single cells according to cell type when using different, commonly used, methods for dimension reduction.* ***A.*** *Using the default Principal Component Analysis.* ***B.*** *Using the default t-SNE with 1000 iterations and 40 perplexity.* ***C.*** *Using the default UMAP.* ***D.*** *Using t-SNE as before but with the Jaccard-Index turned into a distance measure.* ***E.*** *Using the default UMAP method with the Jaccard-Index turned into a distance measure.*

### Supplementary Note 8 – Analyzing the optimal size of the imputations sets

For different imputed sets derived from SIMPA's imputed probabilities the clustering was evaluated in comparison to the imputation set from the reference-free ATAC-seq imputation method (SCALE). The clustering was evaluated using the Davies-Bouldin index (Davies and Bouldin, 1979) based on the results after dimension reduction applied on the imputed or sparse (0 bins added by SIMPA) datasets. The scikit-learn implementation for Python was used to calculate the Davies-Bouldin index (for more details about the dimension reduction procedure and the pathway enrichment analysis we refer to **Supplementary Note 7**).

### Supplementary Note 9 - Implementation details and high-performance computing (HPC)

As described, SIMPA trains a Random Forest classification model specifically for each bin of each single cell. However, we observed that there are bins that have identical class vectors across the target-specific reference experiment set requiring only one model. This reduces the number of classification models that must be trained for one single cell of the H3K4me3 data (5 kb bin size) on average from approximately 314,000 to 138,000.

In order to reduce run time spent on one single cell, SIMPA was implemented using an Open MPI interface for Python (mpi4py 3.0.2). The computationally heavy part of training classification models can be distributed to many CPU cores. We observed high CPU efficiency (> 97%) and 11 GB memory usage when using one full compute node (40 cores, 128 GB RAM, Intel® Xeon® Processor E5-2630 v4) with a runtime of approximately 15 minutes for one single cell. We recommend using SIMPA within a cloud or high-performance computing system. Considering the different validations in this manuscript based on the single-cell profiles for H3K4me3 with 5 kb bins, we applied SIMPA to ~7,500 cases for which 1.2 billion classification models were trained in total. These computations ran on the HPC system MOGON II (JGU, Mainz) in ~24 hours (this can be faster depending on the general workload of the system). Nevertheless, SIMPA can be applied on a standard computer. For one single cell on 5 kb resolution, it takes approximately 120 or 70 minutes using 2 or 4 cores, respectively, of an Intel® Core™ i5-4590 CPU @ 3.30GHz with 8GB RAM.

InterSIMPA trains only one classification model, interprets it, and terminates within seconds.
